# The unique value of zero prediction errors in reinforcement learning

**DOI:** 10.64898/2026.07.13.738284

**Authors:** B. Lloyd, A. Kikumoto, F. Wurm, M.-L. Vives

**Affiliations:** Institute of Psychology, Leiden University, Leiden, Netherlands; Department of Psychology, University of Maryland, College Park

## Abstract

Updating beliefs when necessary is at the cornerstone of learning. A fundamental problem is to describe under which conditions humans update their internal models of the world. The general assumption across animal, human, and artificial learning models^1–3^ has been that updates occur when outcomes deviate from expectations. Perfectly predicted outcomes cause no learning. As a result, no research has examined cases in which predictions are surprisingly perfect. Here, we empirically test this assumption and find that zero prediction errors are psychologically unique. We show that after zero prediction errors, momentary happiness is the highest, and belief updates do indeed occur in a pattern that cannot be reproduced by a benchmark model. We present a new model that captures this non-linear pattern in belief updating by postulating that zero prediction errors elicit a distinct latent belief state, guiding subsequent updating. This latent state then tracks neural activity patterns measured with EEG precisely when zero prediction errors occur, exactly as the model would predict. Crucially, the strength of the neural activity during this time window exhibits a dissociation in predicting the next belief update depending on whether feedback was a zero prediction error or a regular prediction error. Overall, we provide strong evidence that surprisingly perfect predictions are treated in a unique, non-linear fashion at affective, behavioral and neural levels. Being surprisingly accurate can function as a distinct belief updating signal, conforming trial-and-hit learning.

## Introduction

Through experience, we form new beliefs, which drive expectations (“bus arrives at 9 am”). Beliefs are then updated or maintained following an error principle: When expectations are violated (bus arrived at 09:10 am), beliefs are updated. When expectations are met, beliefs are maintained. Learning is thus mostly driven by a mismatch between expectations and outcomes. This basic principle was postulated by Rescorla & Wagner^1^, and has become the basis of most learning models, from simple associations like classical conditioning^1,4,5^ to more complex cognitive processes, such as social interactions^6–9^, financial choices^10,11^, cognitive control^12,13^, and working memory^14^. Sometimes, however, our predictions are *surprisingly* accurate in stochastic outcomes, where there is always a certain degree of uncertainty (bus actually arrived at 9 am). We propose that human cognition is particularly attuned to these surprising zero prediction errors so that, when they occur, affective, behavioral, and neural processes treat them non-linearly, in contrast with what current models assume^1,5^. If that is the case, it would be indicative that perfect, reality-matching expectations constitute, ironically, a special case of error feedback processing.

The impact of zero prediction errors on affect is relatively straightforward to anticipate if we consider that organisms are intrinsically motivated to reduce uncertainty. In humans, a canonical example is the certainty premium: subjective value increases non-linearly when an outcome changes from highly likely (probability = 0.90) to fully certain (probability = 1), suggesting that certainty itself carries value^15^. Similarly, large prediction errors and uncertain outcomes are experienced as aversive^16^, and increases of societal uncertainty are associated with negative affect^17^. This preference for predictability is also present in non-human animals; rats show heightened stress responses when aversive outcomes become unpredictable^18^. At a computational level, active inference accounts propose that organisms are fundamentally driven to minimize prediction error and uncertainty^19,20^ and that a precise match between expectation and outcome signals successful resolution of uncertainty. All considered, zero prediction errors should be intrinsically rewarding, giving rise to positive affect ^21,22^. Interestingly, in current models of momentary happiness ^23,24^, zero prediction errors are treated as a regular instance of prediction errors, reflecting the common, non-special treatment zero prediction errors have received in the literature. Here, we contrast it to an alternative model that weights non-linearly to zero prediction errors. This model can capture the effect that people feel the happiest when their predictions are perfectly matched by reality.

Zero prediction errors are likely to have a positive impact in our affective system: being accurate feels good. A more fundamental question is how people adapt to instances in which outcomes are perfectly predicted. Classical models assume that learning occurs when reality does not match expectations, that is, when there are prediction errors^1,3,4^. Later, more sophisticated accounts show that prediction errors can impact internal representations beyond the general belief (“bus arrives at 9 am”) in, for example, the level of certainty around this belief^25–27^. From another point of view, zero prediction errors may be informative if we consider that belief updating depends also on the uncertainty of the prediction (“I *think* bus comes at 9 am, but I am not sure”). From this perspective, a zero prediction error would carry information about the latent model that generated the prediction, and, by that logic, drive behavioral change during learning.

The main question then is which computational process better captures how people adapt to zero prediction errors. Inspired by recent work showing that large prediction errors trigger the formation of a separate latent belief state^28–30^, we postulate that a similar process follows suit for zero prediction errors. That is, zero prediction errors trigger the creation of a new latent state, which is solely responsible for tracking outcomes that led to zero prediction errors. Following this tenet, we introduce the Zero Prediction Error (ZePE) model, which assumes that people hold two latent states in parallel during learning: one for regular prediction errors, and another one for zero prediction errors, both of which inform the next prediction independently. We contrast this with the current benchmark model^5^, which assumes that learning is driven exclusively by prediction errors, such that larger deviations between expected and observed outcomes produce larger belief updates. Like most models, the benchmark model treats zero prediction errors as an instance where no learning occurs. In ZePE, however, zero prediction errors form a special instance of feedback processing, and, hence, non-linearly treated during learning.

At the neural level, prediction error processing is relatively well characterized: midbrain dopaminergic neurons encode reward prediction errors, increasing firing for better-than-expected outcomes and decreasing firing for worse-than-expected outcomes^3,31–33^. Crucially, when a reward is perfectly predicted, dopaminergic activity remains at baseline, suggesting that zero prediction errors are not represented by a distinct dopaminergic response. This perspective is also reflected in human research using scalp-recorded event-related potentials (ERPs)^34^ to investigate reward prediction error signalling^35,36^, where zero prediction errors are typically treated as the absence of a learning signal rather than as a meaningful computational state. Overall, the frontocentral feedback-related negativity and the subsequent centroparietal P300 have been associated with motivational salience, belief updating, and higher-order evaluation of outcomes^34,37,38^. Importantly, these interpretations are typically grounded in reinforcement learning accounts in which ERP amplitude is expected to vary as a function of reward prediction error^34,36,39^. We challenge this assumption by testing whether zero prediction errors are treated in a non-linear fashion, that is, eliciting larger activity than what would be expected from a linear relationship between prediction error size and neural activity. Preliminary evidence^40^ shows greater neural activity to perfectly predicted outcomes than anticipated, but it was regarded more as a nuisance in the few cases where zero prediction errors randomly occurred rather than a robust empirical finding. As such, those cases were disregarded for further analyses. Here, we provide a direct empirical test by directly manipulating zero prediction errors and link it to the model-base predictions of the new computational model, ZePE.

We provide a first systematic investigation of zero prediction errors as potentially meaningful events in human reinforcement learning. Participants performed a task in which they repeatedly predicted rewards drawn from Gaussian distributions that varied in uncertainty. Crucially, a subset of trials was experimentally constrained such that outcomes exactly matched participants’ predictions, producing true zero prediction errors without participants having explicit knowledge of the manipulation. This design allowed us to examine whether perfectly predicted outcomes influence affective experience, subsequent belief updating, and neural feedback processing.

Our approach was three-fold. First, we tested whether zero prediction errors shape positive affect beyond what can be explained by reward magnitude alone and found that momentary happiness was highest following zero prediction errors. Second, we showed that zero prediction errors influence subsequent belief updating and found that learning was best explained by the ZePE model. Third, we found that zero prediction errors are reflected in canonical electrophysiological signatures of feedback processing in a non-linear manner and that they elicited encodable neural responses linked to the latent state postulated by ZePE. Together, this work challenges the ongoing assumption that no learning occurs when expectations are perfectly met, suggesting that zero prediction errors are a special cue that people use during learning.

## Results

To investigate the impact of zero prediction errors on momentary happiness and learning, participants made trial-by-trial predictions about outcomes generated from a probabilistic reward distribution of varying widths, and were asked “how happy are you at this moment?”^24^ every three trials (see Fig. 1A, adapted from^41^ and^42^). Right after each prediction, participants were presented with the actual reward and the resulting prediction error. Crucially, only outcomes could determine reward; prediction accuracy was not incentivized. Unbeknownst to participants, 15 trials out of 80 in each block were altered such that the reward outcome exactly matched the prediction the participant made, producing a zero prediction error. Data were collected across two experiments (total N = 137; an online experiment, n = 97, and an EEG experiment, n = 40). Behavioral analyses were conducted first for the online experiment and then replicated in the laboratory for the EEG experiment (Supplementary Table 3-5). We present the behavioral results pooled for simplification purposes.

**Figure 1.**
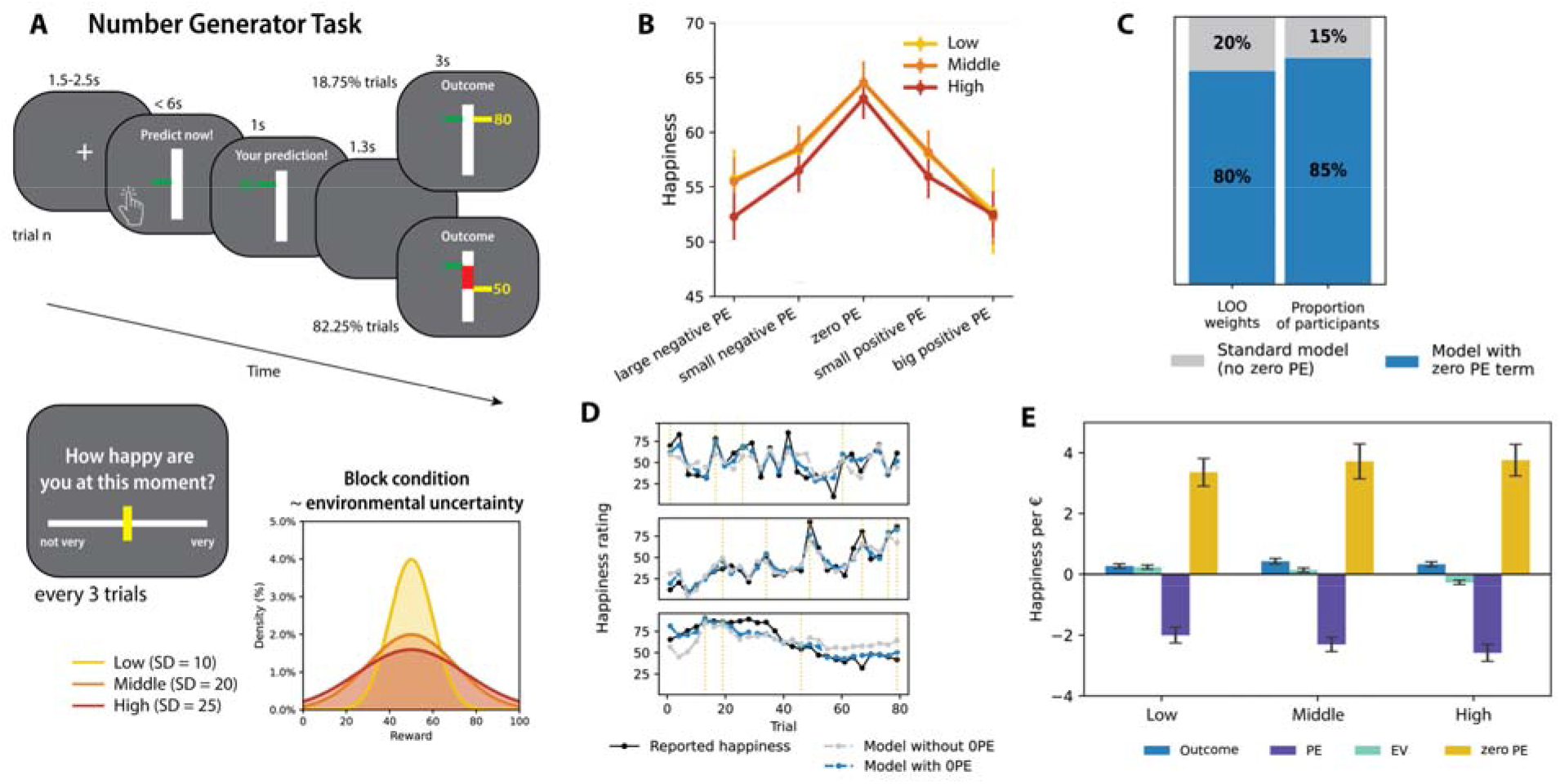
Zero prediction errors elicit the highest happiness. A) Overview of the Number Generator Task. On each trial, participants predicted an upcoming outcome drawn from a Gaussian distribution of varying uncertainty (low, medium, high). After each prediction, the actual outcome and resulting prediction error were shown. Example trials illustrate zero prediction error trials (top) and standard prediction error trials (bottom). Every three trials, participants reported their happiness on a 100-point slider (0 = “not very” to 100 = “very”). B) Happiness showed an inverted-U relationship with prediction error, peaking at zero and decreasing for larger negative and positive prediction errors. Regular negative and positive prediction errors are binned for visualization: small [1–25], large [26–100]. C) Example subjects illustrating model fits. Reported happiness is shown in black, model-predicted happiness from the model including a zero-prediction-error term in blue (dashed), and predictions from the model without this term in gray (dashed). Zero-prediction-error trials are indicated by yellow dashed vertical lines. D) Our computational model adding a zero prediction error term to a computational model^24^ higher predictive accuracy than the original model and received greater model weight, indicating that zero prediction errors contributed uniquely to trial-by-trial happiness. E) Zero prediction errors were the strongest predictor of happiness, with larger posterior coefficient estimates than all other predictors (all *p*_corr_ < .001). Error bars represent SEM. [PE: Prediction error; EV: Expected value].

### People are the happiest after zero prediction errors

We first validated our measure of subjective well-being (see Supplementary Information), to then explore the relationship between prediction error magnitude and momentary happiness. Previous findings suggest that happiness scales together with the size of positive prediction errors, that is, the larger the unanticipated positive reward, the happier people feel^23,24^. This was not replicated in our task: mixed-effects regression revealed that the magnitude of prediction errors on the previous trial did not linearly predict happiness (*b* = 0.044, 95% CI [−0.02, 0.11], *z* = 1.27, *p* = .20). In contrast, we hypothesized that people value perfect predictions, and hence, feel the happiest after a zero prediction error, and less happy as prediction error magnitude increases, either positively or negatively. This inverted-U-shaped pattern was confirmed in a mixed-effects regression predicting happiness ratings from continuous prediction error values, including both linear and quadratic terms. The quadratic term was significant and negative, consistent with lower happiness as prediction errors deviated further from zero (*b* = −0.002, 95% CI [−0.003, −0.001], *z* = −3.21, *p* = .001): Happiness peaked at zero prediction errors and declined for both large negative and large positive prediction errors (see Fig. 1C). Participants were happier after perfectly predicting an outcome than after receiving a larger monetary reward than expected. In our task, certainty had higher intrinsic value than surprising positive rewards. Participants were also sensitive to reward magnitude, as happiness was significantly higher after larger outcomes (*b* = 0.73, 95% CI [0.330, 1.136], *z* = 3.56, *p* < .001). In fact, the inverted-U relationship between prediction error and happiness was slightly stronger at higher reward levels (see Supplementary Table 1), suggesting that zero prediction errors had the greatest affective benefit when the reward itself was high.

Finally, we extended a computational model of momentary well-being^24^ (Methods for more details below) by including a unique free parameter for zero prediction error trials. The new zero prediction error model—compared to the original model—substantially improved fit (ΔELPD = 541.86, SE = 61.55), had higher predictive accuracy and greater model weight (0.80 vs. 0.20; Fig. 1D). Posterior estimated parameters indicated zero prediction errors as the strongest contribution to happiness (mean = 3.61, 95% CI [3.20, 4.14]; Fig. 1E; see Supplementary Table 2 for posterior means across levels of uncertainty), with previously described components^24^ of much lesser relevance (Outcome: mean = 0.34, 95% CI [0.14, 0.53] and Prediction Error: mean = −2.30, 95% CI [−2.50, −2.08]). During learning, happiness is non-linearly driven by perfect predictions.

### Zero prediction errors drive a separate belief state: the ZePE model

How do people react to zero prediction errors during learning? To test this, we computed the average absolute prediction updates after zero prediction errors. Against the predictions of the Pearce-Hall model^5^, considered the benchmark in this task^41,42^ (see Fig. 2A), we found a pronounced bump around zero prediction error, with larger updates on and around zero prediction error trials. We then tested the ZePE model, inspired by recent work showing that large prediction errors can trigger the formation of a separate latent belief state^30^. In this framework, surprising outcomes can give rise to a new state that is tracked in parallel to ongoing learning. Extending this idea, we hypothesized that the surprising nature of zero prediction errors in stochastic environments may similarly give rise to a distinct latent state. According to the ZePE model, participants are assumed to maintain two latent beliefs about the outcome distribution. One belief is updated using standard prediction errors according to the Pearce–Hall learning rule, capturing gradual belief updating following mismatches between expected and observed outcomes. The second belief specifically tracks trials in which the prediction exactly matches the outcome (zero prediction errors), reflecting a state in which the participant’s current expectation is perfectly accurate. Behavior is then generated from a weighted combination of these two belief states, controlled by a free parameter, ω, which determines the relative influence of the zero prediction error belief state and the standard prediction error driven belief state on subsequent predictions. Values of ω closer to 1 indicate that predictions are mostly driven by the zero prediction error belief state, whereas values closer to 0 indicate that predictions are primarily driven by the standard prediction error driven belief state.

**Figure 2.**
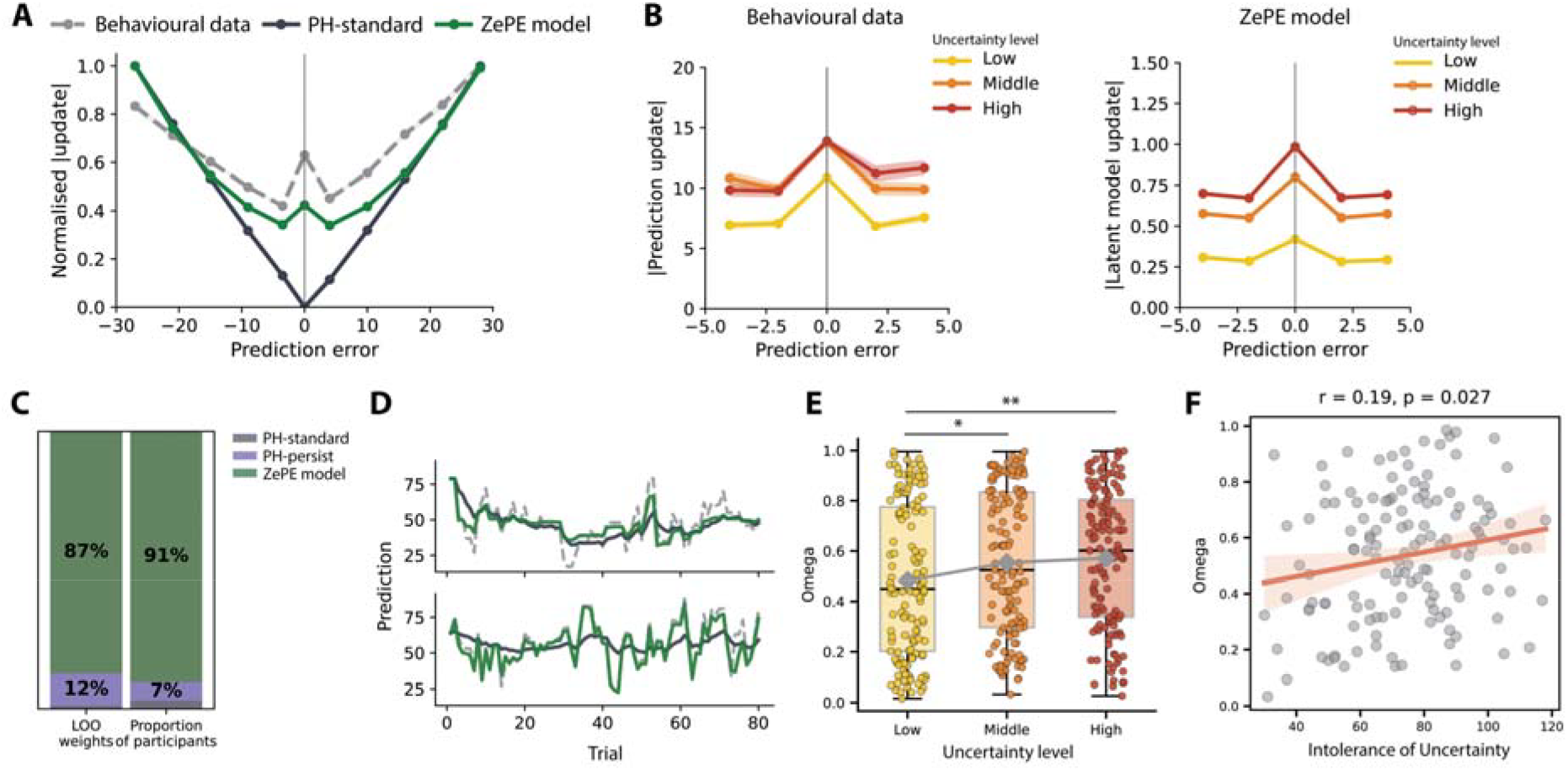
ZePE captures the qualitative pattern of larger updates than expected after zero prediction errors. A) Behavioural data showed a W-shaped pattern centred on zero prediction error, which was not predicted by the benchmark PH-standard model but was predicted by ZePE. B) Looking more closely at small prediction errors, updating after zero prediction error increased under high uncertainty in both the behavioural data and the ZePE model simulations. C) The ZePE model provided the best fit relative to the benchmark PH-standard model and the PH-persist model, based on both LOO weights and the proportion of participants best fit by each model, indicating that zero prediction errors play an important role in learning. Model fits were collapsed across tasks for visualisation purposes only. D) Example participant showing trial-by-trial fit between observed behavioral predictions and latent belief estimates from the PH-standard and ZePE models. E) Estimates of the free parameter ω increased with uncertainty, differing between Low and Middle and Low and High levels of uncertainty. F) Participants with higher intolerance of uncertainty showed larger ω values, indicating that individuals high in uncertainty aversion place more weight on zero prediction erros during learning. Data points depict individual participants and error bars and shaded regions denote SEM. (*: *p* < .05; **: *p* < .005).

ZePE could reproduce the novel qualitative pattern, showing enhanced latent updating around zero prediction error (see Fig. 2A). Furthermore, we then examined the impact of environmental uncertainty in this W pattern. Results revealed that the pattern changed, showing a larger update for higher uncertainty relative to lower uncertainty (Middle: *b* = 1.12, 95% CI [0.418, 1.821], *z* = 3.13, *p* = .002; High: *b* = 1.52, 95% CI [0.825, 2.215], *z* = 4.29, *p* < .001; see Fig. 2B [left]). ZePE reproduced this pattern, showing enhanced latent updating around zero prediction error across uncertainty levels. The influence of zero prediction errors became more pronounced under higher environmental uncertainty (Fig. 2B [right]). Together, these qualitative patterns provide behavioral support for the ZePE model by showing that participants updated not only in response to classical prediction errors, but also around zero prediction error events.

We then tested if ZePE was also the best fitting model, as the qualitative patterns would suggest, by using Bayesian hierarchical modelling (separate sequences for each participant and block), to get more stable parameter estimates, and compared using leave-one-out cross-validation (LOO), which estimates each model’s out-of-sample predictive accuracy^43,44^. We contrasted ZePE with the benchmark, Pearce-Hall model (PH-standard model)^5^ and an alternative model that gives a persistent effects to zero prediction errors (PH-persist model, see details in the Method section and Supplementary Information). All models showed excellent model- and parameter-recovery performance (see Supplementary Information). In both experiments, model comparison strongly favored the ZePE model (ELPD_LOO online_ = −93,468.97, ELPD_LOO EEG_ = −67,533.91), which substantially outperformed both the PH-persist model (ΔELPD_LOO online_ = 1,109.92, SE = 77.25 ΔELPD_LOO EEG_ = 1,551.16, SE = 68.37) and the PH-standard model (ΔELPD ΔELPD_LOO online_ = 4,806.30, SE = 116.80, ΔELPD_LOO EEG_ = 5,062.62, SE = 108.72). Across the two experiments (see Fig. 2C,D), the ZePE model received most of the LOO stacking weight (0.83 online, 0.92 EEG), compared with PH-persist model (0.15 online, 0.08 EEG), and for the standard PH-standard model (0.02 online, 0.0003 EEG). At the participant level, the ZePE model provided the best predictive fit for most individuals (84 participants out of 97 online, all participants in EEG). Together, these results indicate that learning is best captured by a model in which zero prediction errors are non-linearly weighted, inducing a new distinct belief state that guides updating.

### Omega captures the impact of environmental uncertainty and individual differences

An interesting, novel pattern observed is that zero prediction errors elicit a larger belief update under higher environmental uncertainty, providing evidence that surprise modulates the weight people give to zero prediction errors in their updating. To further interrogate this phenomenon at the computational level, we examined which parameter of ZePE was sensitive to changes of environment uncertainty. A repeated-measures ANOVA revealed a significant effect of uncertainty on the new free-parameter, ω, which arbitrates how much weight is given to each latent state (*F*(2,268) = 6.59, *p* = .002, 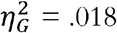). Pairwise comparisons showed that ω was significantly higher in High compared to Low blocks (*t*(134) = −3.41, *p*_corr_= .003, *d*_z_ = −0.29) and in Middle compared to Low blocks (*t*(134) = −2.55, *p*_corr_ = .024, *d*_z_ = −0.22), while there was no difference between Middle and High blocks (*t*(134) = 0.77, *p*_corr_ = .444, *d*_z_ = 0.07; Fig. 2E). This pattern indicates that ω increases with environmental uncertainty. When the environment becomes less predictable, people rely more on surprisingly perfect predictions. Additional simulations varying ω revealed corresponding patterns in behavioral updating (see Supplementary Information).

One important use of computational models has been to provide a better characterization of which cognitive processes are associated with certain individual differences (e.g., anxiety, depression) by analysing its relationship with models’ free parameters^45–47^. Individuals’ levels of intolerance of uncertainty may be especially relevant to zero prediction errors because individuals averse to uncertainty differ in how they interpret and resolve probabilistic outcomes^48^. We therefore examined whether individuals’ uncertainty attitudes^49^ predicted individual differences in ω, while including environmental uncertainty as a within-subject factor. This analysis revealed a significant main effect of intolerance of uncertainty on ω (*b* = 0.002, 95% CI [0.0003,0.0041], *p* = 0.03; Fig. 2G), indicating that individuals who are more intolerant of uncertainty tended to place greater weight on the zero prediction error latent belief state (see Supplementary Information for the full regression). These results suggest reliance on zero prediction errors as captured by the ZePE free-parameter, omega, can capture relevant variance at the individual level, which opens the avenue for future studies investigating other constructs, like psychosis^50^.

### Zero prediction errors are non-linearly weighted at the neural level

We first examined EEG activity traditionally associated with prediction error processing. Absolute prediction error was associated with an early, broadly distributed frontocentral–parietal component approximately 200–320 ms after outcome onset, followed by a later, spatially widespread component approximately 420–700 ms after outcome onset (see Supplementary Information for full results). These temporal and spatial profiles are broadly consistent with canonical reward-related components related to the P300, with the earlier positivity resembling a frontocentral P3a-like response and the later, more extended component resembling a centroparietal P3b-like response ^51^.

To characterize how zero prediction errors relate to neural error signals, we examined nuisance-corrected EEG responses across five conditions: large negative, small negative, zero, small positive, large (Fig. 3A). Based on previous work showing that outcome-locked surprise is reflected in a central P300-like positivity spanning approximately 300–700 ms and moving from frontocentral to centro-parietal electrodes, we focused our primary ERP analysis on Fz, Cz, and Pz within a late positive time window ^52^. In general, all prediction error conditions showed a similar temporal profile, characterized by a negative early component (~100–350 ms) followed by a late positive component (~400–900 ms). Notably, zero prediction errors were associated with an enhanced late positive response that was comparable in magnitude to that observed for large positive prediction errors, while differing from other prediction error conditions, which showed reduced late activity. This effect was particularly evident when averaging activity within the 500–800 ms window (Fig. 3B), where responses differed significantly across prediction error categories, *F*(4,156) = 12.9, *p* < .001, 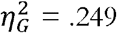. Follow-up paired comparisons, FDR-corrected across all pairwise tests, showed that zero prediction errors elicited larger late positive responses than large negative (*p*_corr_ < .001, *d*_z_ = 0.65), small negative (*p*_corr_ < .001, *d*_z_ = 0.85), and small positive prediction errors (*p*_corr_ < .001, *d*_z_ = 0.60), but did not differ significantly from large positive prediction errors, (*p*_corr_ = .395, *d*_z_ = −0.15). Together, these results reveal a non-linear relationship between prediction error magnitude and EEG activity, with maximal responses for zero and large positive prediction errors. Zero prediction error are non-linearly treated at the neural level as well.

**Figure 3.**
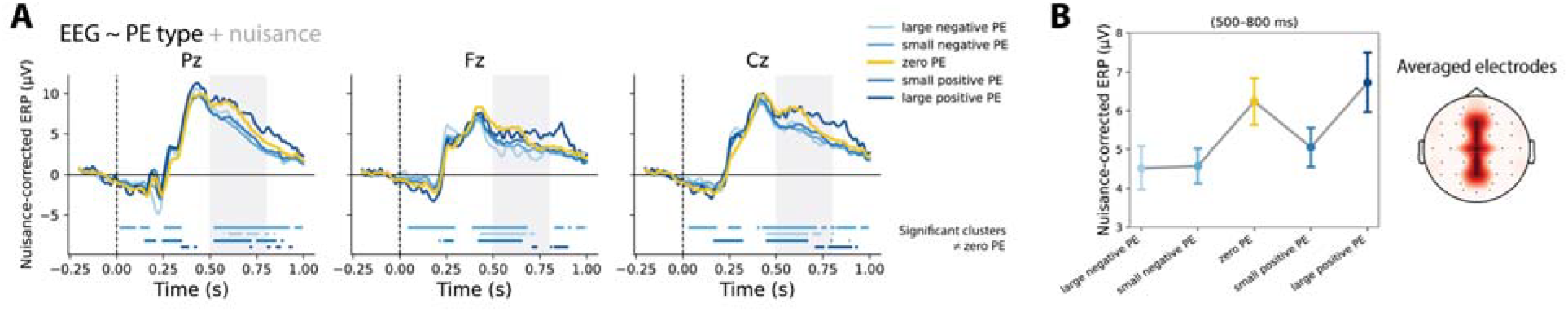
Electrophysiological signatures of zero prediction errors. A) Patterns of EEG responses following zero prediction errors resemble those of large positive reward prediction errors. B) Average EEG responses varied nonlinearly with prediction error. Nuisance-corrected ERPs at Pz, Fz, and Cz are shown for binned prediction errors (large negative to large positive; zero prediction error as reference). The shaded window (500–800 ms) corresponds to the averaged values shown in panel B averaged across electrodes Pz, Fz, and Cz. Horizontal bars indicate spatiotemporal clusters that survived cluster-based permutation testing (*p* < .05). Error bars denote SEM. [PE: Prediction error].

### Neural encoding of the zero prediction error latent state predicts unique belief updates

In line with the premise that zero prediction errors are uniquely treated, we find a higher amplitude than expected in nuisance corrected ERPs. Our new computational model, ZePE, provides another, more precise prediction in how the brain reacts to zero predictions errors. Specifically, it predicts that

EEG activity would reflect the dissociative nature of the ZePE model. To test this, in each trial, we computed the difference between the observed outcome and zero prediction error latent belief state on one side, 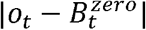, and the standard prediction error latent belief state on the other side, 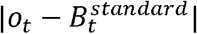. Importantly, we then entered both latent states, together with their interactions with trial type, into a timepoint × channel generalized linear model: 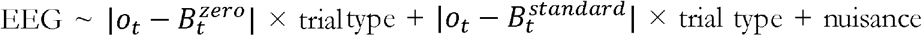 regressors. Trial type was defined as 0 when trial belongs to standard prediction error and 1 when trial belongs to zero prediction error. This allowed us to test whether EEG activity uniquely tracked deviations from each latent belief state. The main effect of 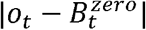 yielded no significant clusters on standard trials (all *p* > .05). In addition, we observed a significant interaction between 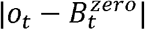 and trial type from approximately 500-1000 ms after outcome onset (*p* = .002; *t* = 2.87), indicating that this latent-state prediction error signal was larger for zero prediction error trials than standard prediction error trials. By contrast, the main effect of 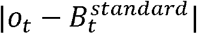 yielded a significant positive cluster spanning approximately 355–1,000 ms after outcome onset (*p* = .027, *t* = 2.61), whereas no significant clusters were observed for the interaction between 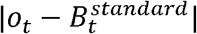 and trial type (all *p* > .05). To explore the direction of these trial-type-specific effects, we examined condition specific slopes derived from the interaction terms (Fig. 4A). These simple-slope analyses should be interpreted cautiously, particularly where they rely on comparing significant and non-significant effects rather than a direct interaction contrast^53^. With this caveat, 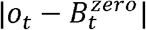 showed a significant positive cluster on zero prediction error trials, spanning approximately 500-900 ms (*p* = .003; *t* = 2.78), whereas no significant clusters were observed on standard trials (all *p* > .05). These results suggest that EEG activity tracks distinct latent-state deviations derived from ZePE’s model, distinguishing zero prediction and regular prediction errors.

**Figure 4.**
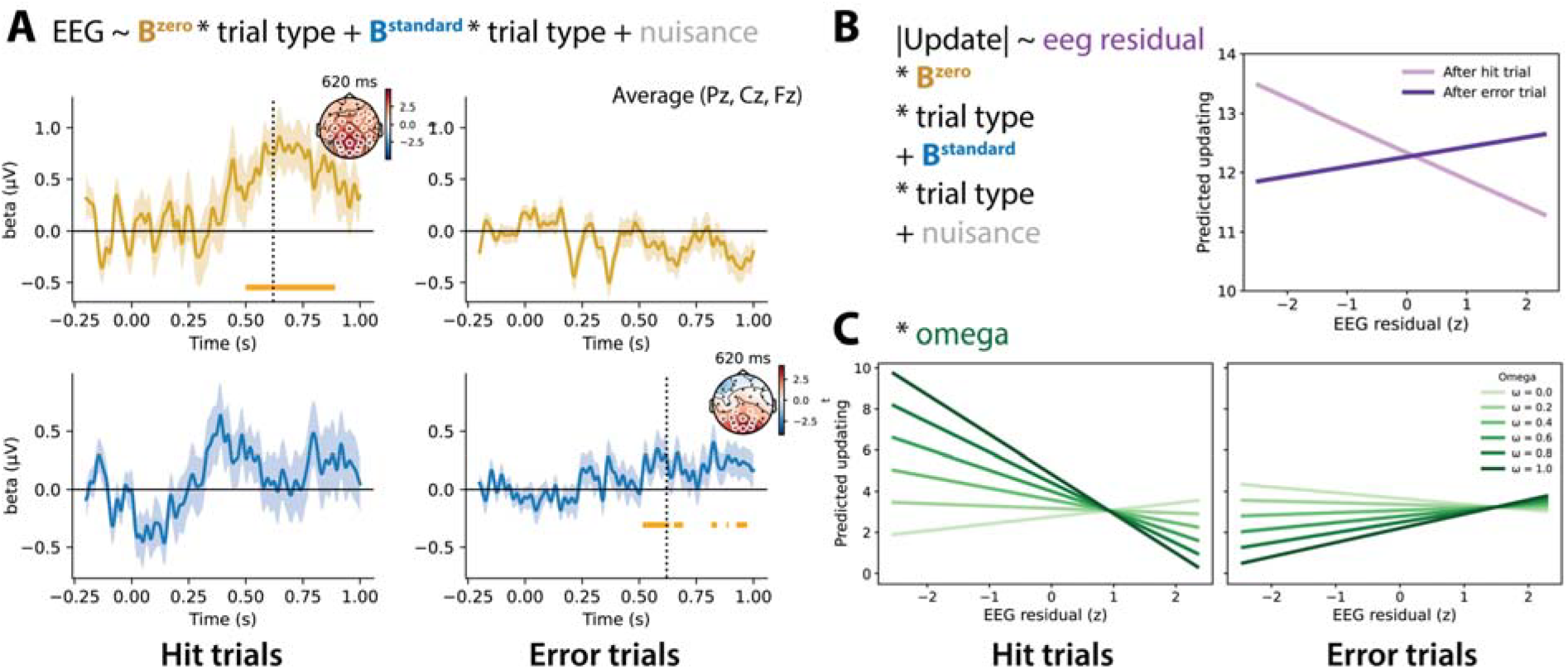
Residual EEG activity predicts subsequent belief updating differently for zero prediction error and standard trials. A) EEG activity tracked prediction errors relative to the model-derived zero and standard latent belief states differently across trial types. Time courses are averaged across frontocentral electrodes (Pz, Fz, Cz), with insets showing topographies at peak latency (620 ms). B) PPI analysis revealed residual EEG activity predicted subsequent updating in opposite directions across trial types: larger updates after standard trials, but reduced updates after zero prediction error trials. C) This trial-type-dependent relationship was further modulated by ω. Horizontal bars and white circles indicate spatiotemporal clusters that survived cluster-based permutation testing (*p* < .05). Error bars and shaded regions denote SEM. (in figure B^*Zero*^refers to 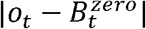 and B^*Standard*^ refers to 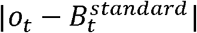; PE: Prediction error).

The conclusion that latent states derived by the ZePE model explained unique variance in feedback-related EEG activity would be further strengthened by evidence that residual EEG activity associated with these signals also predicted subsequent behavioral updating. To test this, we carried out a psychophysical interaction analysis (PPI). First, we extracted trial-wise EEG residuals from the previous mass-univariate model, reflecting feedback-related neural activity not explained by the latent belief-state predictors, trial type, or nuisance regressors. We then tested whether these residual EEG signals predicted absolute belief updating on the following trial. Absolute prediction updates on trial *n+1* were modeled as a function of the model-derived latent deviations 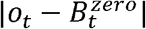 and 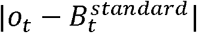, trial type, and their interactions, while controlling for block uncertainty and trial number. Crucially, the model also included interactions between the EEG residuals, the latent belief-state predictors, and trial type, allowing us to test whether residual feedback-related EEG activity moderated the relationship between latent computational signals and subsequent behavioral updating. We found that the relationship between residual EEG activity and subsequent updating differed as a function of trial type. Greater residual EEG activity was associated with reduced subsequent updating following zero prediction error trials, *b* = −0.46, 95% CI [−0.85, −0.07], *p* = .020. This relationship was significantly more positive during standard prediction error trials, as indicated by a residual EEG × trial type interaction, *b* = 0.63, 95% CI [0.19, 1.06], *p* = .005 (Fig. 4B; Supplementary Table 6). Thus, stronger residual neural responses predicted attenuated updating following perfect predictions, but enhanced updating following standard learning trials, indicating a neural dissociation between zero prediction errors and regular prediction errors.

Finally, we examined whether the relationship between residual EEG activity and subsequent behavioral updating was further modulated by the weighting parameter ω from the ZePE model. To this end, we extended the previous PPI analysis by including ω and its interactions with residual EEG activity, trial type, and the latent belief-state signals in the multilevel model predicting absolute prediction updates on trial *n+1*. Critically, we observed a significant interaction between residual EEG activity, trial type, and ω, *b* = 3.24, 95% CI ([1.48,4.99], *z* = 3.61, *p* < .001 (Fig. 4C; Supplementary Table 7). Specifically, the more participants weighted zero prediction errors trials (higher ω values) in their predictions, the larger the dissociation between trial types: residual EEG activity was increasingly associated with attenuated updating following zero prediction error trials, but enhanced updating following standard prediction error trials. The new free parameter of ZePE, omega, arbitrates the neural dissociation of error feedback processing in belief updating.

## Discussion

Our findings challenge the longstanding assumption that perfectly predicted outcomes are psychologically neutral. Zero prediction errors consistently emerged as unique events, treated non-linearly at affective, behavioural, and neural levels. First, the highest levels of momentary happiness were reported after zero prediction errors, uncovering a novel, inverted-U pattern with prediction error magnitude. In fact, zero prediction errors elicited more happiness than obtaining a reward higher than expected and had the strongest influence in happiness. Second, we uncovered a new qualitative pattern in belief updating that was not predicted by the benchmark model. Updates were larger than anticipated after zero and small prediction errors, following a W-shape, with a bump precisely for zero prediction errors. A new model, ZePE, captured this pattern by postulating two latent states, one for regular prediction errors and another one for zero prediction errors. The influence of this state increased under higher environmental uncertainty, suggesting that perfect predictions are weighted more when they become more surprising. Finally, electrophysiological analyses revealed that zero prediction errors elicited robust and sustained outcome-locked EEG responses in a non-linear fashion, and selectively engaged neural signals linked to the latent zero prediction error belief state. Critically, residual EEG activity predicted subsequent belief updating in opposite directions depending on trial type: greater neural responses were associated with attenuated updating following zero prediction errors, but enhanced updating following standard prediction error trials, a result that was further modulated by the weight each participant gave to each latent state during learning. Together, these findings show that zero prediction errors uniquely shape affective experience, learning, and neural processing of feedback.

We provide the first evidence that participants consistently reported the highest momentary happiness following zero prediction errors. A central implication of this finding is that perfectly predicted outcomes are intrinsically rewarding. Importantly, prediction accuracy was not incentivized: extra reward was determined only by the outcome. Zero prediction errors may be rewarding because they minimize the discrepancy between the organism’s current internal state and a desired goal state of successful prediction^56^, signalling that the environment has been successfully captured. This interpretation resonates with work on moments of insight, where sudden gains in understanding are often accompanied by happiness and positive affect^57,58^. Such moments are also characterized by a reduction in uncertainty^59^. Thus, the happiness caused by zero prediction errors may reflect a positive internal signal linked to the formation of an accurate model of the world.

A major implication of these findings concerns the computation of surprise during learning. Although prediction errors are typically treated as the primary learning signal^5,60,61^, the present findings suggest that the absence of prediction error can also become informative when it is itself unexpected. The role of surprise for latent state creation is present in Bayesian and uncertainty-sensitive accounts of learning ^25,27,62–64^, and distributional reinforcement-learning models^65–67^. Surprise, however, is directly linked to the size of the prediction error. Here, we generated a new computational model, ZePE, in which we postulate that surprise can also occur for zero prediction errors (and probably small prediction errors as well, as suggested by the W shape discovered), and lead to the creation of two latent states. In stable environments, zero prediction error events may be adequately handled by a single latent state. By contrast, in dynamic decision-making environments, where exact prediction is unlikely, surprisingly small predictions can provide a signal, from an information-theory point of view, about the world. In line with this, we found that the weight participants gave to zero prediction errors, captured by omega, increased together with environmental uncertainty, when zero prediction errors were indeed more surprising. Future research could explore the degree to which this applies as well for small prediction errors and if people use them as a learning signal as well.

The present findings raise important questions about the neural systems that encode prediction errors in general and zero prediction errors in particular. First, we found that zero prediction errors elicited as much amplitude as large positive prediction errors, which casts doubts on the classical interpretation of EEG signal scaling only on prediction error magnitude^35,39^, confirming previous, unexpected patterns^40^. Furthermore, we uncovered a neural dissociation in how EEG activity within the same time-window predicted belief updating. This dissociation was further modulated by ZePE’s new free-parameter, omega, such that the more people relied on zero prediction errors during learning, the larger the difference in how the neural activity predicted belief update. An interesting possibility is that omega is shaped by dopaminergic mechanisms, given the central role of dopamine in reward prediction error signaling^3,31^ and belief updating^68^. Locus coeruleus–noradrenaline function also tracks changes in learning rate in volatile environments, while acetylcholine and noradrenaline have been proposed to support different forms of uncertainty processing^27,69–71^. Omega thus may index a neuromodulatory tuning process that determines whether neural activity promotes stabilizing or updating beliefs, which would reshape our understanding of neural reward processing.

Establishing the main laws of learning is one of the most pressing questions in society, as non-human agents become more sophisticated using basic principles like trial-and-error. As established by Thorndike’s law of effect: behaviors followed by satisfying consequences are more likely to be repeated, while behaviors followed by unpleasant consequences become less likely to occur. Here, we re-orient a field overly focused on trial-and-error learning, usually linked to unpleasant consequences, towards learning from surprisingly accurate predictions, usually linked to pleasant consequences, conforming trial-and-hit learning.

## Methods (STAR)

### Online experiment

#### Participants

One hundred and one individuals carried out the number generation task. Four participants were removed due to no variance in their happiness ratings, leaving a final sample of 97 participants (48 females, 47 males, 2 NA) between the ages of 18 and 45 in Experiment 1. All participants were recruited from the United States via Prolific, spoke fluent English, and provided written informed consent. The study was approved by the Psychology Research Ethics Committee of Leiden University (CEP: M.L. Vives Moya-V1-5664).

#### Experimental Task

Participants performed a number generation task designed to assess learning of reward distribution means. The task was programmed in PsychoPy, hosted online via Pavlovia, and distributed through Prolific. All participants completed the study remotely on their own desktop computers. Each participant completed three blocks of 80 trials. In each block, outcomes were drawn from a Gaussian distribution that varied in both precision (standard deviation [SD]: low = 10, medium = 20, high = 25; block order counterbalanced across participants) and expected value (mean randomly drawn from 40 to 60). Each trial consisted of several phases. First, a prediction screen appeared, displaying a vertical bar and a movable slider (range: 0–100). Participants used the mouse to indicate their prediction of the upcoming outcome. After confirming their response, the selected value remained on screen for 1200 ms, followed by the presentation of an abstract image for 1500 ms. This image presentation was included to address a separate research question beyond the scope of this article. The trial concluded with feedback (see Fig. 1A), displaying: the actual outcome (randomly sampled from the current distribution), the participant’s prediction, and a visual representation of the prediction error (defined as outcome – prediction). Participants were told that different lottery machines generated outcomes clustered around particular numbers, but were not informed about the underlying uncertainty level or distribution width associated with each block. In each block, 15 trials (18.75%) were randomly selected to produce a zero PE (i.e., the outcome exactly matched the participant’s prediction). Participants were not informed about the inclusion of these trials. These zero prediction error trials were randomly distributed throughout each block and were imposed independently of the Gaussian reward distribution. Every three trials, participants were asked to report their current mood by answering: “How happy are you right now?” via a slider scale (0-100), following the method introduced by ^24^. One trial was randomly selected at the end of the experiment to determine bonus pay.

#### Procedure

At the start of the experiment, participants were told they would be interacting with three different “lottery machines”. They were told that although outcomes may appear random, each machine tended to generate rewards closer to specific values, and their goal was to learn these underlying tendencies across trials. Participants were also informed that one trial from each block would be randomly selected, and the outcome from each of those three trials would be converted into cents, summed across blocks, and paid out as a bonus at the end of the experiment. Prior to the main task, participants completed five practice trials. Following the lottery task, they filled out basic demographic information and completed the 27-item intolerance of uncertainty scale ^49^. The experiment took approximately 75 minutes to complete, and participants were compensated at a rate of €7.15 per hour. Participants who failed either of two attention checks—one embedded in the task instructions and one embedded at the end of the intolerance of uncertainty scale—were excluded from analysis and not compensated.

### EEG experiment

#### Participants

40 participants (27 females, 12 males, 1 NA), aged between 18 and 50, participated in Experiment 2. All participants were recruited through SONA, tested in person at Leiden University, spoke fluent English, and provided written informed consent. Two participants were missing two blocks due to time constraints, and one block from another participant was excluded because the participant fell asleep.

#### Experimental Task

The same task was used as in the online experiment, with the following modifications. First, the outcome range was increased from 0–100 to 0–200 and only one trial was selected to determine the bonus payment, which was converted directly into euros (e.g., 125 points = €12.50). Second, the task consisted of six blocks instead of three. For each block, the distribution mean was randomly drawn from either a low (75–99) or high (100–125) outcome range. Each uncertainty level (Low, Middle, High) was presented twice, and each outcome range appeared three times, with block order counterbalanced across participants. Third, the abstract image presented after the prediction phase in the online experiment was omitted.

#### Procedure

Participants entered the EEG laboratory, and the setup procedure lasted approximately 20 minutes. EEG and pupil size were recorded throughout the experiment. The entire session lasted approximately two hours, and participants were compensated €22 for their time.

#### EEG acquisition

EEG was recorded using a BioSemi ActiveTwo system (BioSemi) with 32 Ag–AgCl electrodes arranged according to the international 10–20 system. To monitor ocular activity, additional electrodes were placed above and below the left eye (vertical electrooculogram [EOG]) and on the outer canthi of both eyes (horizontal EOG). Data were continuously recorded at 512 Hz. Electrode impedance was kept within acceptable limits according to BioSemi’s active electrode system specifications.

#### EEG preprocessing

EEG analyses were conducted using the MNE-Python package ^72^. The data were first re-referenced to the average of the left and right mastoid electrodes, and then band-pass filtered between 1 Hz and 30 Hz using a FIR filter with a ‘firwin’ design. Independent component analysis (ICA) was used to remove artifacts from the EEG data. First, we decomposed the continuous EEG signal into 15 components using the Infomax algorithm implemented in MNE. Next, we applied automatic component classification using MNE’s *label_components* function, which labels components based on their spatial and temporal characteristics. Components associated with ocular artifacts (e.g., blinks), cardiac signals, and muscle activity were automatically identified and excluded from the data.

The cleaned signal was then segmented into epochs time-locked to outcome onset, spanning from – 200 ms to +1000 ms. These epochs were baseline-corrected by subtracting the mean voltage in the 200 ms window prior to outcome onset. Epochs containing large residual artifacts were excluded from further analysis using a peak-to-peak amplitude threshold of 250 µV. A total of 52 bad epochs were dropped across all participants, leaving 99.07% clean trials for analysis.

#### EEG analysis

EEG data were analyzed using a mass univariate regression approach. For each participant, EEG amplitude at every electrode and timepoint relative to outcome onset was regressed onto a design matrix containing task-relevant regressors of interest alongside nuisance covariates. Regressors of interest varied across models and included trial-wise estimates of absolute prediction error (z-scored), trial type (0 = standard trial, 1 = zero prediction error trial), and model-derived latent prediction error signals (e.g., 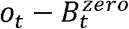 and 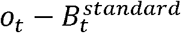), as well as their interactions with trial type (standard trials as reference). Nuisance regressors included uncertainty condition (Low, Middle, High), block (1 to 6), and trial number within block. This approach yielded regression coefficients (*b*) at each electrode and timepoint for each participant. Group-level statistics were performed using one-sample *t*-tests on these coefficients across participants, resulting in spatiotemporal maps of *t*-statistics for each regressor of interest. To correct for multiple comparisons, we employed a cluster-based permutation test ^73,74^. Spatiotemporal clusters were defined as contiguous samples across electrodes and timepoints exceeding a cluster-forming threshold (two-tailed, *p* < .05). Cluster-level statistics were computed by summing *t*-values within each cluster. The significance of observed clusters was assessed against a null distribution generated via 2000 permutations using sign-flipping across participants. Clusters with *p*-values below α = .05 were considered significant.

### Intolerance of Uncertainty

Uncertainty attitudes were assessed using the 27-item Intolerance of Uncertainty Scale ^49^. Participants indicated their agreement with each statement on a Likert scale (e.g., 1 = not at all characteristic of me to 5 = entirely characteristic of me). Item scores were summed to yield a total IUS score, with higher values reflecting greater intolerance of uncertainty.

#### Behavioral analyses

We used the Python function ‘mixedlm’ (package: statsmodels) to estimate mixed-effects linear regression models ^75^. Except for the analyses involving the EEG signal, data from both studies were pooled to maximize statistical power. Our analyses were based on a set of core model structures, with specific predictors or interaction terms added where relevant to the hypothesis tested and reported in the Results section.

First, we modeled participants’ affect ratings as a function of prediction error and uncertainty level. To examine whether affect varied nonlinearly with prediction error, we included both linear signed prediction error (PE) and quadratic prediction error (PE^2^) terms. Second, we modeled trial-by-trial absolute prediction updating as a function of absolute prediction error, zero prediction error status (coded 0/1), and uncertainty level (Low, Middle, High). Interactions between prediction error, zero-prediction-error status, and uncertainty were included where relevant to test whether the shape of the prediction-error–update relationship differed across trial types and uncertainty levels. Mixed-effects models included participant-specific random intercepts and slopes, with trial number and block number entered as regressors of no interest. To examine whether the ZePE mixture parameter σ varied with environmental uncertainty, we conducted a repeated-measures analysis of variance with uncertainty level as a within-participant factor. Significant effects were followed by pairwise comparisons between uncertainty levels. We additionally used a mixed-effects regression to test whether intolerance of uncertainty predicted σ, while accounting for environmental uncertainty as a within-participant factor. Data from the EEG experiment, which were collected on a 0–200 scale, were normalized to a 0–100 scale to match the online experiment. All statistical tests were two-sided with an alpha level of .05. Effect sizes were reported as regression coefficients (*b*) for regression and mixed-effects models, generalized eta squared 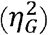 for analyses of variance, and paired-samples Cohen’s *d*_z_ for within-participant comparisons. Cohen’s *d*_z_ was calculated as the mean within-participant difference divided by the standard deviation of the difference scores. Unless otherwise stated, data are reported as mean ± SEM, and corrected p-values, denoted *p*_corr_, refer to false discovery rate correction ^76^.

### Computational modeling

#### Happiness model

We modeled happiness using a modified version of the computational model proposed by Rutledge et al. (2014), which includes terms for outcome value (O), expected value (*Ev)*, absolute prediction error (|*PE*|), and zero prediction error (*zero PE*). Here, o_*j*_ denotes the reward outcome on trial *j*; *EV*_*j*_ denotes the expected value on trial *j*; |*EV*_*j*_ | denotes the absolute prediction error; and *zeroPE*_*j*_ is a binary indicator coding whether trial *j* produced a zero prediction error.

The original model did not include the *zero PE* term and was defined as follows:

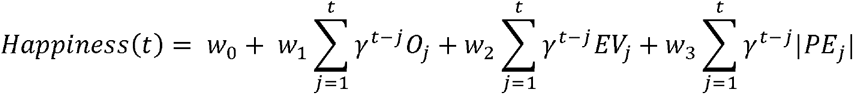

All continuous predictors (*O,Ev*, (|*PE*|) were z-scored within each participant and block to ensure comparability of regression weights across predictors and to account for differences in scaling, whereas the *zero PE* term was coded as a binary variable (0 = standard PE, 1 = zero PE). According to the modified model, happiness at trial *t* is defined as:

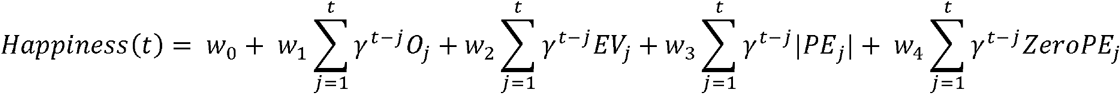

where *t* is trial number, *w*_*0*_ is a constant, and *w*_1-4_ are weights that capture the influence of each factor. The impact of past events decays exponentially over time according to the forgetting factor γ (0 ≤ γ ≤1), such that more recent outcomes have a greater influence on current happiness.

#### Learning models

To capture how learning unfolds trial-by-trial, we fitted several reinforcement learning models to participants’ predictions, initially inspired by Haarsma et al. (2021). On each trial *t*, PE was defined as the discrepancy between the observed outcome *o*_*t*_ and the participant’s prediction *y*_*t*_:

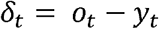

In all models, the observed predictions were assumed to be generated from a latent belief state B_*t*_ with some Gaussian noise:

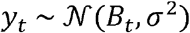

Across the models, learning was governed by a Pearce-Hall-style dynamic learning rate α _*t*_, which increased with recent unsigned prediction error:

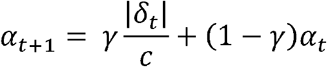

where *c* is the scaling constant (here *c* = 100) and *γ* ∈ [0,1] determines the extent to which the recent surprise influences the current learning rate. Learning rate was constrained to the interval [0,1]. Models differed in the rule governing the evolution of *B*_*t*_. Parameters were estimated using hierarchical Bayesian modeling implemented in PyMC ^77^, with parameters estimated for each participant and block (see model fitting section for details).

##### Pearce-Hall (PH) model

In the standard PH-standard model, latent expectations were updated according to the current PE, weighted by the dynamic learning rate term:

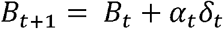

Therefore, larger learning rates meant predictions shifted more strongly toward the current outcome. The free parameters of this model were the initial learning rate *α*_0_, the Pearce-Hall weighting parameter *γ* and the observation noise parameter *σ*, which captures the amount of noise or variability in participants’ predictions.

##### PH-persist model

The PH-persist model extended the standard PH-standard model by allowing additional updating on zero prediction error trials indexed by *z*_*t*_, where *z*_*t*_ = 1 on zero PE trials and *z*_*t*_ = 0 otherwise. As in the standard PH-standard model, latent beliefs were first updated according to:

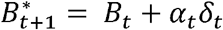

On zero prediction error trials (*z*_*t*_=1), the updated belief was then shifted further toward the current outcome according to:

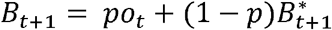

where *o*_*t*_ is the reward outcome and *p* is a persistence parameter (bounded by the interval ∈ [0,2]) controlling the extent to which the updated belief is pulled toward the current outcome. Equivalently,

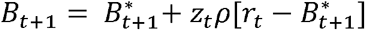

when *z*_*t*_ = 0, the model reduces to the standard PH update. When *z*_*t*_ = 1, the updated belief is pulled further toward the current outcome. The free parameters were *α*_0_, *γ,ρ*, and *σ*.

##### ZePE model

The ZePE model assumed that predictions were generated from a weighted mixture of two latent beliefs, one associated with zero prediction error trials, 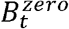, and one associated with standard (non-zero) prediction error trials, 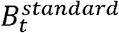. Both latent beliefs evolved according to the same dynamic Pearce-Hall learning rate *α*_*t*_, but were updated selectively depending on trial type.

First, candidate beliefs were computed for both states:

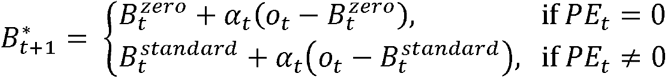

Then, on zero prediction error trials (*PE*_*t*_ = 0), only the zero prediction error state was updated, while the standard prediction error state remained unchanged. Conversely, on standard prediction error trials (*PE*_*t*_ ≠ 0), only the standard prediction error belief was updated.

Therefore, the model maintains separate latent beliefs for zero prediction error and standard prediction error events and combines them into an overall prediction. The model prediction on each trial was given by

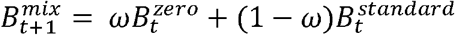

where *ω* ∈ [0,1] determines the relative contribution of the zero prediction error and standard prediction error latent belief states for that trial. The free parameters were *α*_0_, *γ, ω*, and *σ*.

##### Model fitting

Model parameters were estimated using hierarchical Bayesian inference implemented in Python with the PyMC library ^77^. For each model, subject-level parameters were drawn from group-level distributions, allowing partial pooling across participants while preserving individual differences ^43^. Models were fitted separately for the online and EEG experiments, such that partial pooling occurred within each experiment and group-level estimates were experiment specific. Subject-specific parameters were estimated separately for each level of environmental uncertainty. Parameters constrained to the unit interval, including *α*_0_ and *γ*, and, where applicable, *ρ* and *ω* were estimated on the unconstrained real line using hierarchical normal distributions. For each uncertainty level, subject-level raw parameters were drawn from a group-level normal distribution, with normal priors on the group means and half-normal priors on the group standard deviations. Raw parameters were then mapped to the interval (0,1) using a logistic transformation. Observation-noise parameters were modeled hierarchically on the log scale and exponentiated to ensure positivity. Group-level parameters defined the population means and standard deviations of the corresponding subject-level distributions. Posterior inference was performed using Markov chain Monte Carlo sampling with the No-U-Turn Sampler implemented in PyMC. For each model, we ran four independent chains with 1,000 tuning iterations followed by 1,000 sampling iterations per chain. The target acceptance rate was set to 0.95 to improve sampling stability. Convergence was assessed using standard diagnostics, including inspection of trace plots and the potential scale reduction statistic 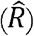. Trial-level log-likelihood values were retained to enable model comparison using leave-one-out cross-validation.

##### Simulation analysis

To visualize the behavioral consequences of the fitted learning models, we performed a simulation analysis using the empirical group-mean parameter estimates obtained from the hierarchical Bayesian model fits. We simulated trial-by-trial predictions and latent belief states from the standard Pearce–Hall model and the ZePE model. For the PH-standard model, simulations used *α*_0_ = 0.17, *γ* = 0.06, and *σ* = 18. For the ZePE model, simulations used *α*_0_ = 0.72, *γ* = 0.21, *ω* = 0.52, and *σ* = 13. For each model, we generated 1,000 simulated blocks for each level of environmental uncertainty, with feedback standard deviations of 10, 20, and 25. Each block contained 80 trials, with zero prediction error trials occurring on 18.75% of trials, matching the empirical task design. For each simulation, the environmental mean was sampled uniformly between 40 and 60 and was held constant across models, allowing direct comparison of the two models under matched task conditions. We then extracted signed and absolute prediction errors, observed prediction updates, and latent belief-state updates from the simulated trial sequences. Observed updates were defined as the change in sampled prediction from trial t to trial *t+1*, whereas latent updates were defined as the corresponding change in the model’s internal belief state.

##### Parameter recovery

To assess whether model parameters could be reliably recovered from behavioral data, we conducted parameter recovery analyses for the three models considered in the main analyses (PH, PH-persist, and ZePE model). For each model, simulated datasets were generated using the same task structure as in the online experiment (240 trials per dataset). On each simulation, model parameters were sampled from uniform distributions spanning the parameter ranges used during model fitting (e.g., *α*_0_, *γ* ∈[ 0,1], *ρ* ∈ [0,2], *ω* ∈[0,1], and *σ* ∈[1,12]). Outcome values were sampled from a Gaussian distribution with mean 60 and standard deviation 20, matching the approximate distribution of lottery outcomes in the task. In the simulation, 18.75% of trials were randomly selected to produce a zero prediction error, paralleling the behavioral study. Using these parameters, we simulated trial-by-trial predictions according to the generative model. Simulated behavioral data were then refit using the same model and fitting procedure as used for the empirical data. This procedure was repeated for 300 simulated datasets per model. Parameter recovery was assessed by computing the correlation between the true generative parameters and the parameters recovered by the fitting procedure. In addition, we computed the bias (mean difference between recovered and true parameters) and root-mean-square error (RMSE) for each parameter. Results of the parameter recovery analyses indicated that the key model parameters (*α*_0_, *γ*, and *σ*, as well as ρ for PH-persist and *ω* for the ZePE model) were reliably recovered across simulations (see Supplementary Information).

##### Model recovery

To assess whether the candidate models could be dissociated, we conducted a model recovery analysis for the three models entered into the main comparison (PH, PH-persist, and ZePE model; Wilson & Collins, 2019). Synthetic datasets were generated from each model using parameter values sampled from the admissible parameter ranges and the same task structure as in the experiment (240 trials per dataset). In total, 200 datasets were simulated, split evenly across the three generative models. Each simulated dataset was then fit with all candidate models, and model identity was recovered using the Bayesian Information Criterion (BIC). Recovery performance was summarized using a confusion matrix showing the probability of selecting each fit model conditional on the true generative model, *p*(fit | simulated), as well as the corresponding inversion matrix, *p* (simulated | fit). The row-normalized confusion matrix quantified recoverability, that is, how often data generated by a given model were best fit by that same model. The column-normalized inversion matrix quantified reliability, that is, how often selection of a given model reflected that model as the true data-generating process. Data generated from each of the three models were correctly identified in a minimum of 92.5% of cases (see Supplementary Information for full results). Together, these analyses assessed whether the PH, PH-persist, and ZePE models were sufficiently distinguishable to support subsequent model comparison on the empirical data.

## Data and code availability

The raw data supporting the findings of this study are publicly available on Figshare: https://figshare.com/projects/Dissociable_neural_encoding_of_zero_prediction_errors_predicts_opposite/276021. The analysis code is available on GitHub: https://github.com/bethlloyd/Zero_prediction_errors.

## Acknowledgements

This work was supported by a Dutch Starter Grant (Startersbeurs) awarded to Marc-Lluís Vives by Leiden University. We would also like to thank Mirjami Tuurna and Cyd Vaandrager for their help with data collection.

