## Supplementary Information for "The unique value of zero prediction errors in reinforcement learning"

**Happiness measure tracks task variables**

We first analyzed whether it was sensitive to aspects of the task. First, in line with previous findings ^1^, subjective well-being decreased over the course of the experiment (mean ± SEM; initial 65 ± 25, final 54 ± 28, *t*(134) = 4.16, *p* < .001; Supplementary Fig. 1A). Furthermore, subjective well-being also captured the negative affect caused by higher levels of uncertainty ^2^, as happiness was lower when rewards were more difficult to predict (Low vs. Middle: b = −1.16, p < .001; Low vs. High: b = −2.90, p < .001; Supplementary Fig. 1B). Together, these results show that our happiness measure can capture expected fluctuations of subjective well-being.

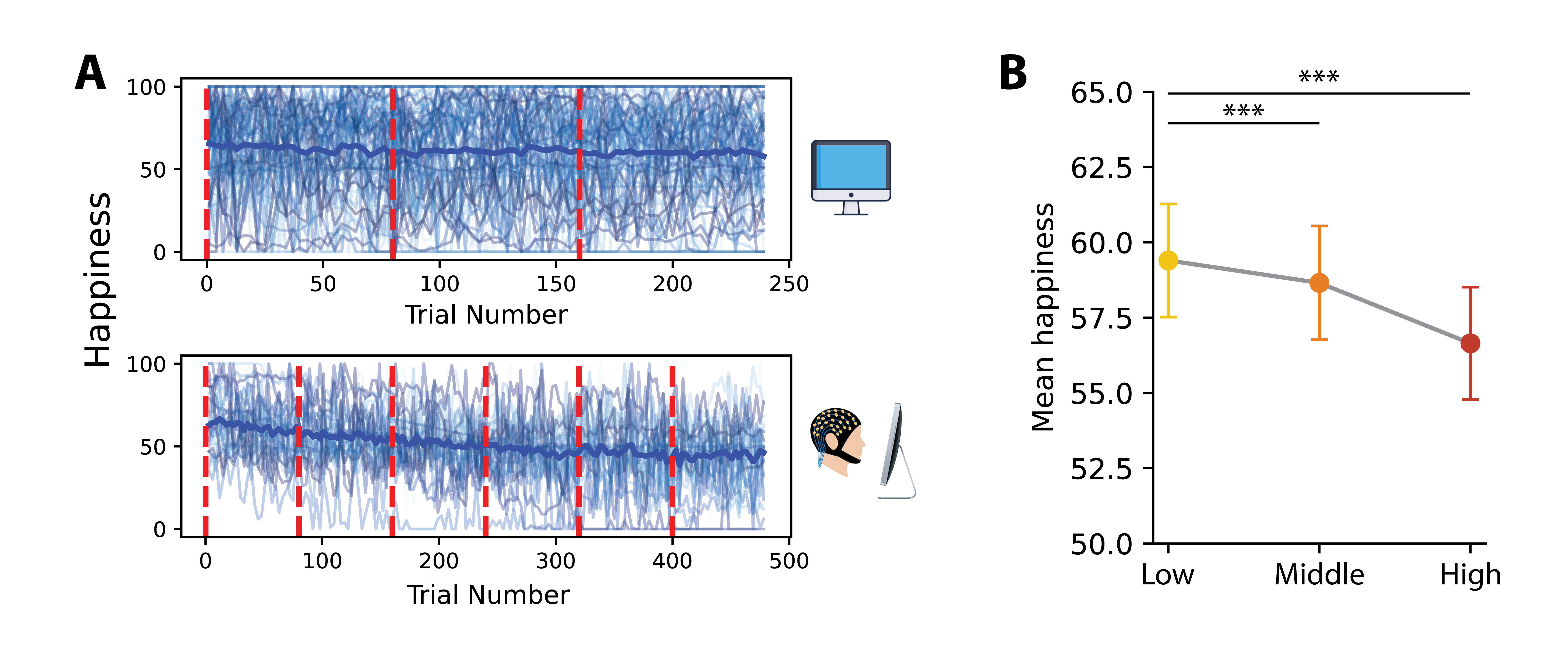

**Supplementary Figure 1. Happiness measure tracks time-on-task and environmental uncertainty**. A) Happiness across the task significantly decreased for the Online and EEG experiments. Red dashed lines refer to the beginning of each block. B) Environmental uncertainty also decreased happiness significantly. (***: *p* < .001).

**Supplementary Table 1.**

*Linear mixed-effects model predicting happiness ratings from prediction error, reward outcome, block, and trial number*

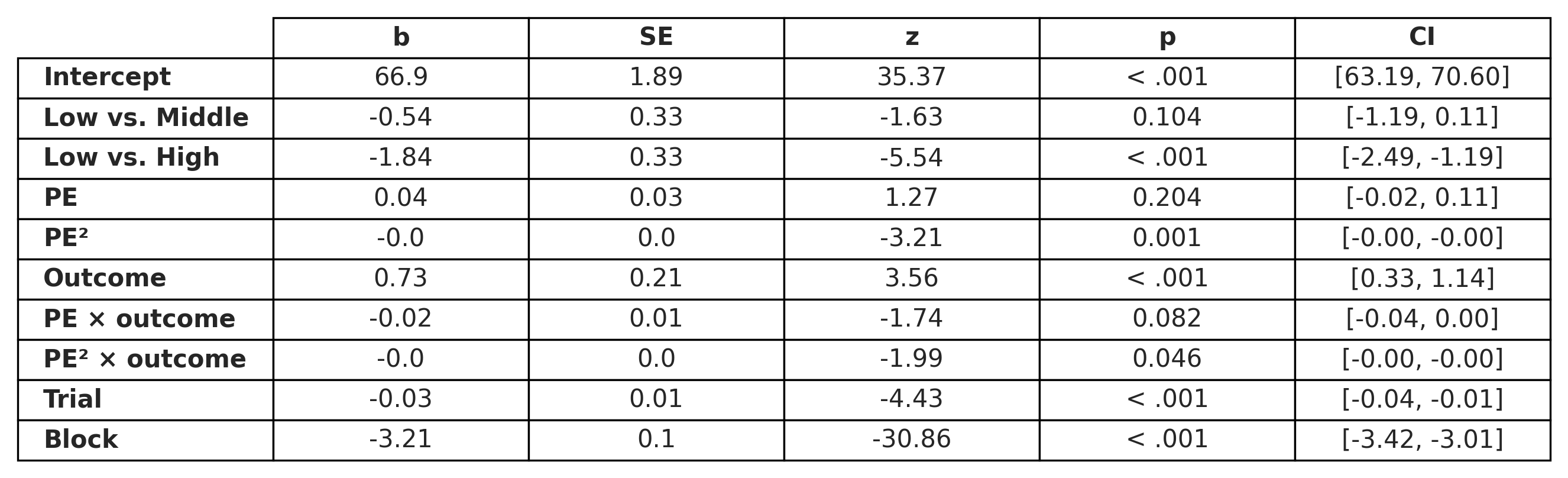

*Note.* Fixed-effect estimates (*b*), standard errors (SE), z-values, p-values, and 95% confidence intervals (CI) are reported. Task uncertainty (standard deviation) was coded with the Low condition as the reference level; coefficients for middle and high therefore reflect contrasts relative to Low.

**Supplementary Table 2.**

*Posterior means (SD) of model parameters across uncertainty levels*

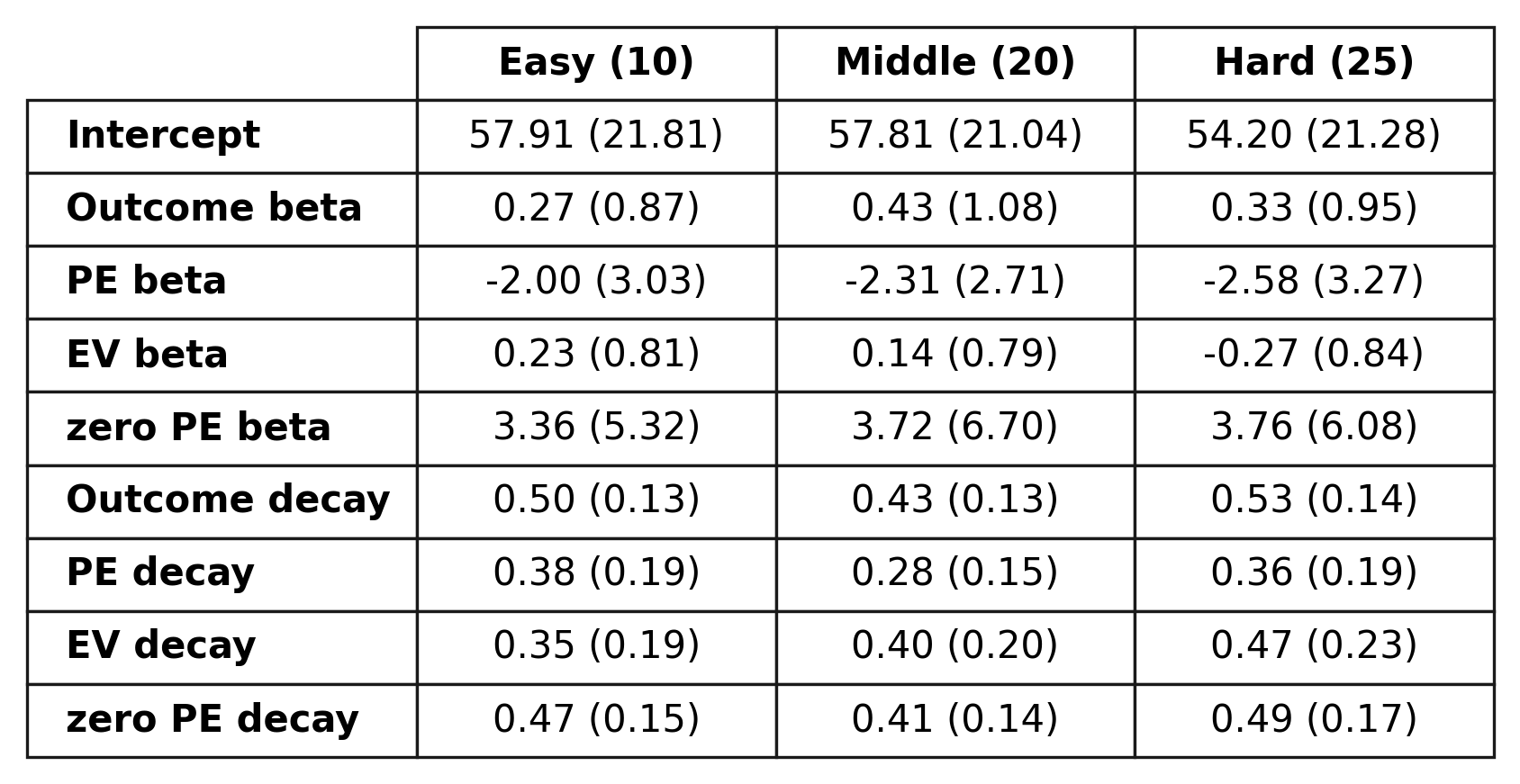

*Note*. Table shows posterior means with standard deviations, estimated from the hierarchical Bayesian happiness model including the zero-prediction error predictor. Parameters are reported separately for each uncerainty level (Low = 10, Middle = 20, High = 25). Outcome, prediction error (PE), expected value (EV), and zero prediction error (zero PE) correspond to regression coefficients predicting momentary happiness. Decay parameters (τ) reflect the temporal integration of each predictor. The intercept reflects baseline happiness. Higher values indicate stronger contributions of each predictor to reported happiness.

**Supplementary Table 3.**

*Task-adjusted combined results: momentary happiness*

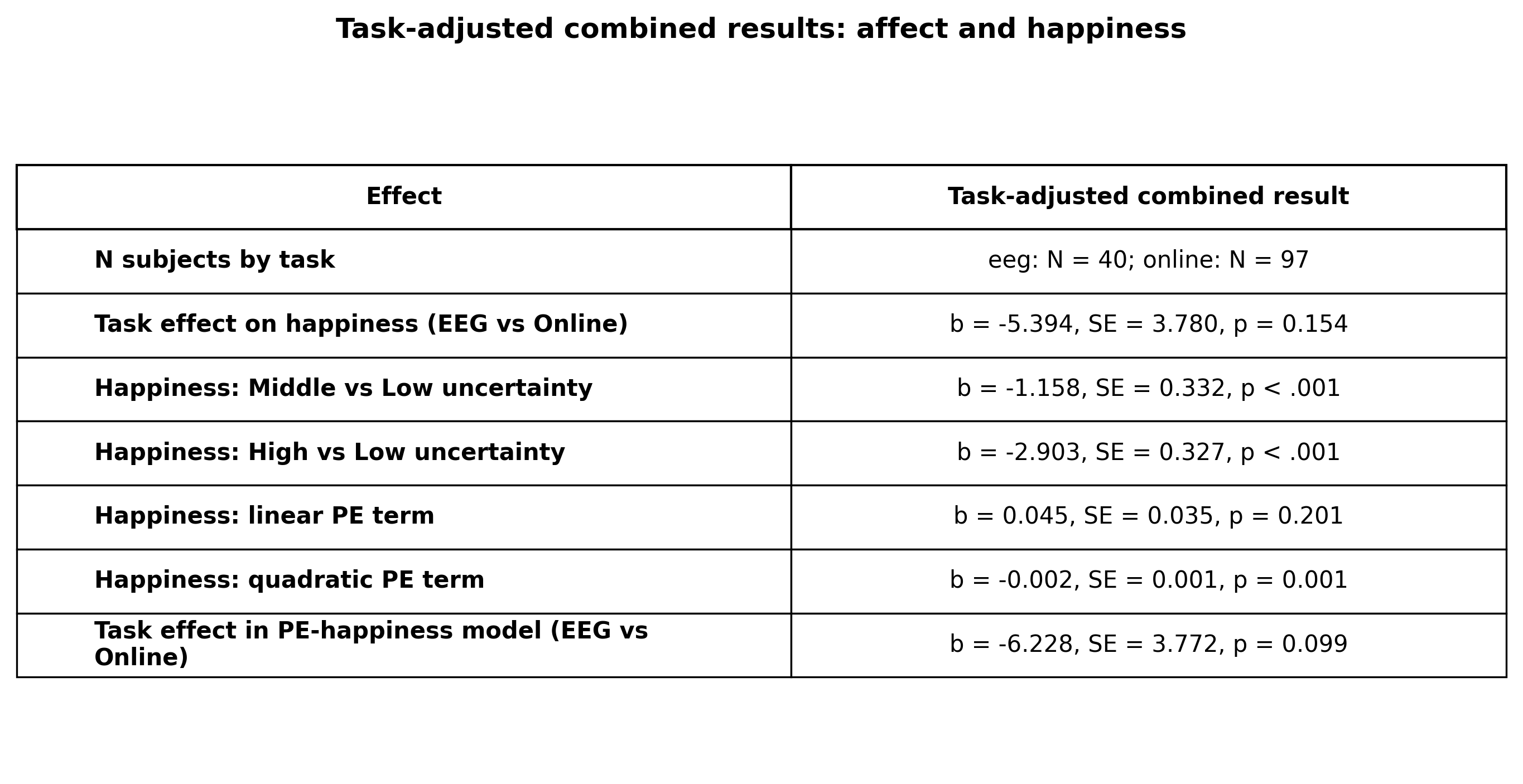

*Note.* Table shows task-adjusted combined results for affect and happiness across the Online and EEG samples. Task was included as an additional fixed-effect regressor, with the online sample as the reference category, to control for dataset differences. Estimates are regression coefficients with standard errors. Happiness decreased with increasing uncertainty, and the quadratic prediction-error effect remained significant after controlling for task, block, trial number, and reward magnitude.

**Supplementary Table 4.**

*Task-adjusted combined results: behavioural learning and updating*

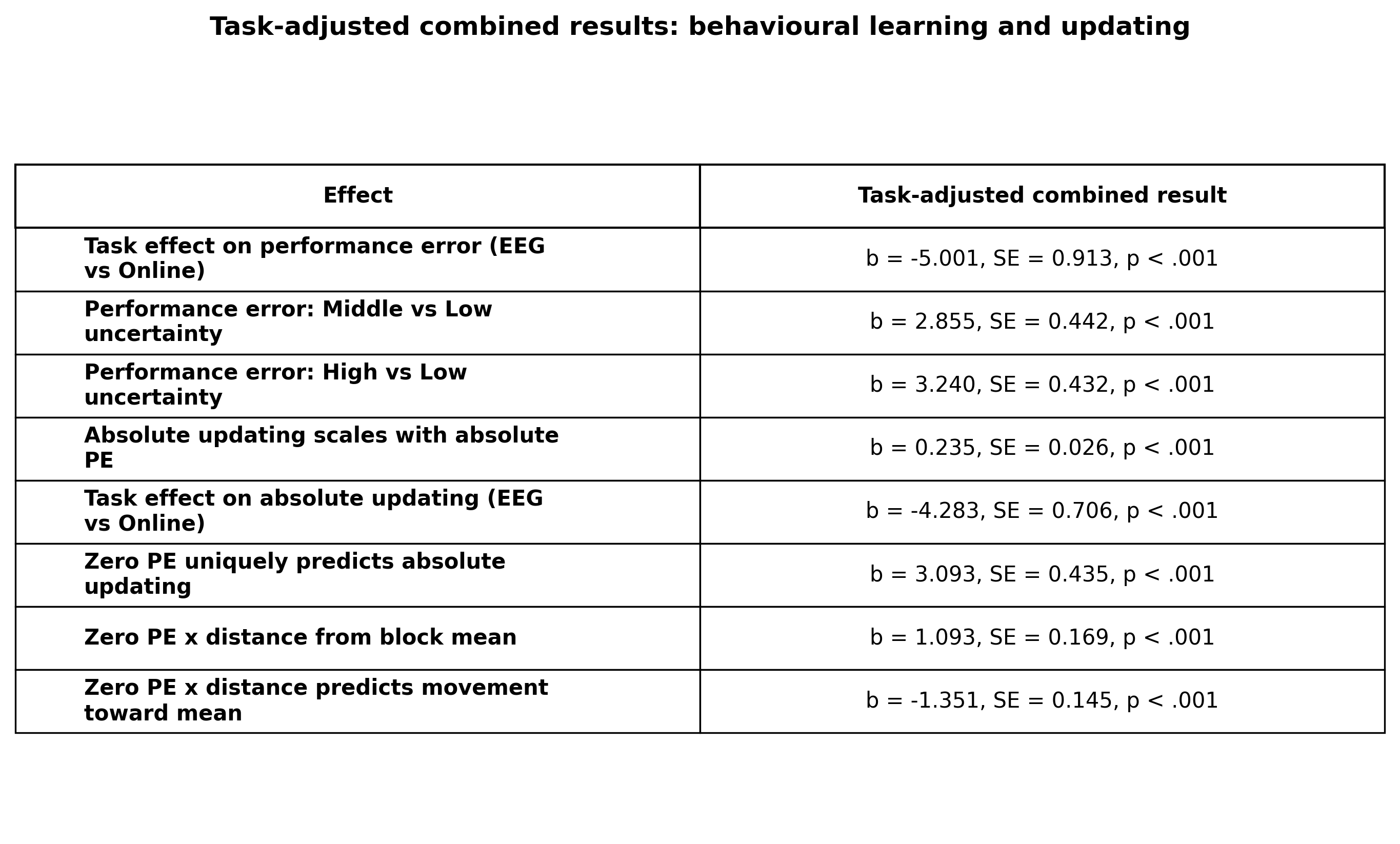

*Note.* Table shows task-adjusted combined results for behavioural learning and updating across the Online and EEG samples. Task was included as an additional regressor in each model to account for differences between datasets, with the online sample as the reference category. The table reports the main effects of uncertainty, absolute prediction error, zero prediction error, and distance from the block mean after controlling for task, block, and trial number. Estimates are regression coefficients with standard errors. These analyses show that the key behavioural effects remained reliable after accounting for task differences.

**PH-persist model: persistent effects of zero prediction errors**

In addition to the benchmark PH-standard model, we also considered an extension of this model: the PH-persist model. In the PH-persist model, receiving a zero prediction error induces a persistent state that influences subsequent learning. Within reinforcement learning frameworks, reduced updating or ‘stickiness’ can emerge either through low learning rates, which limit the influence of new information on existing beliefs, or through an additional preservation parameter that biases behavior toward previous responses ^3,4^. Building on this idea, we hypothesized that the unique nature of zero prediction errors—both in terms of their subjective impact and their role in learning—may dynamically engage such persistence. Specifically, moments of perfect prediction may trigger a temporary bias toward maintaining current beliefs, effectively ‘locking in’ the perfect outcome. In the PH-persist model, perfect predictions therefore modify the learning process beyond the immediate trial, allowing zero prediction errors to shape future belief updates by biasing subsequent predictions toward the confirmed outcome. The degree of persistence is controlled by a free parameter, ρ, which determines how strongly zero prediction errors influence future predictions. Higher values of ρ result in stronger persistence toward the confirmed outcome, whereas lower values of ρ revert to the standard PH update with little to no persistence.

**Model Validation**

We conducted parameter and model recovery analyses to verify that parameters were recoverable and that the candidate models could be reliably distinguished. As shown in Supplementary Fig. 2A, for the PH-standard model, parameter recovery was strong for the Pearce–Hall weighting parameter ($\gamma$, $r=.88$) and the observation-noise parameter ($\sigma$, $r=.96$), while recovery of the initial learning-rate parameter ($\alpha_{0}$) was more modest ($r=.55$) For the **PH-persist model**, all parameters were highly recoverable: ($\rho$, $r=.99$), the Pearce–Hall parameter ($\gamma$, $r=.97$), and the noise parameter ($\sigma$, $r=.99$). Recovery of the initial learning-rate parameter was weaker ($r=.48$), consistent with the limited influence of this parameter beyond the earliest trials. Finally, for the **ZePE model**, recovery was strong for the mixture parameter ($\omega$, $r=.95$) and the noise parameter ($\sigma$, $r=.996$), and good for the Pearce–Hall weighting parameter ($\gamma$, $r=.81$). Recovery of the initial learning-rate parameter was relatively weak ($r=.27$), again reflecting the limited contribution of this parameter after the initial trials of the task. Overall, the parameters central to the theoretical predictions of the models ($\gamma$, $\rho$, and $\omega$) showed high recoverability, indicating that the models could reliably recover the parameters of primary interest. Inspection of the recovery plots confirmed that recovered parameters tracked the true generative parameters without strong parameter trade-offs.

Overall, the models were well recovered (Supplementary Fig. 2B). Data generated from the **PH-standard model** were correctly identified in 92.5% of cases (62/67), with the remaining datasets primarily misclassified as the PH-persist model (7.5%). Similarly, datasets generated from the **PH-persist model** were correctly recovered in 92.5% of cases (62/67), with the remaining 7.5% classified as the PH-standard model (See Supplementary Fig. 2B). For the **ZePE model**, recovery accuracy was 86.4% (57/66). Misclassifications occurred exclusively toward the PH-standard model (13.6%), with no cases in which ZePE model datasets were best fit by the PH-persist model. The inversion matrix further indicated that when the **ZePE model** was selected as the best-fitting model, it always corresponded to data generated from that model (100% reliability). Overall, these results indicate that the candidate models were sufficiently distinguishable to support model comparison on the empirical data.

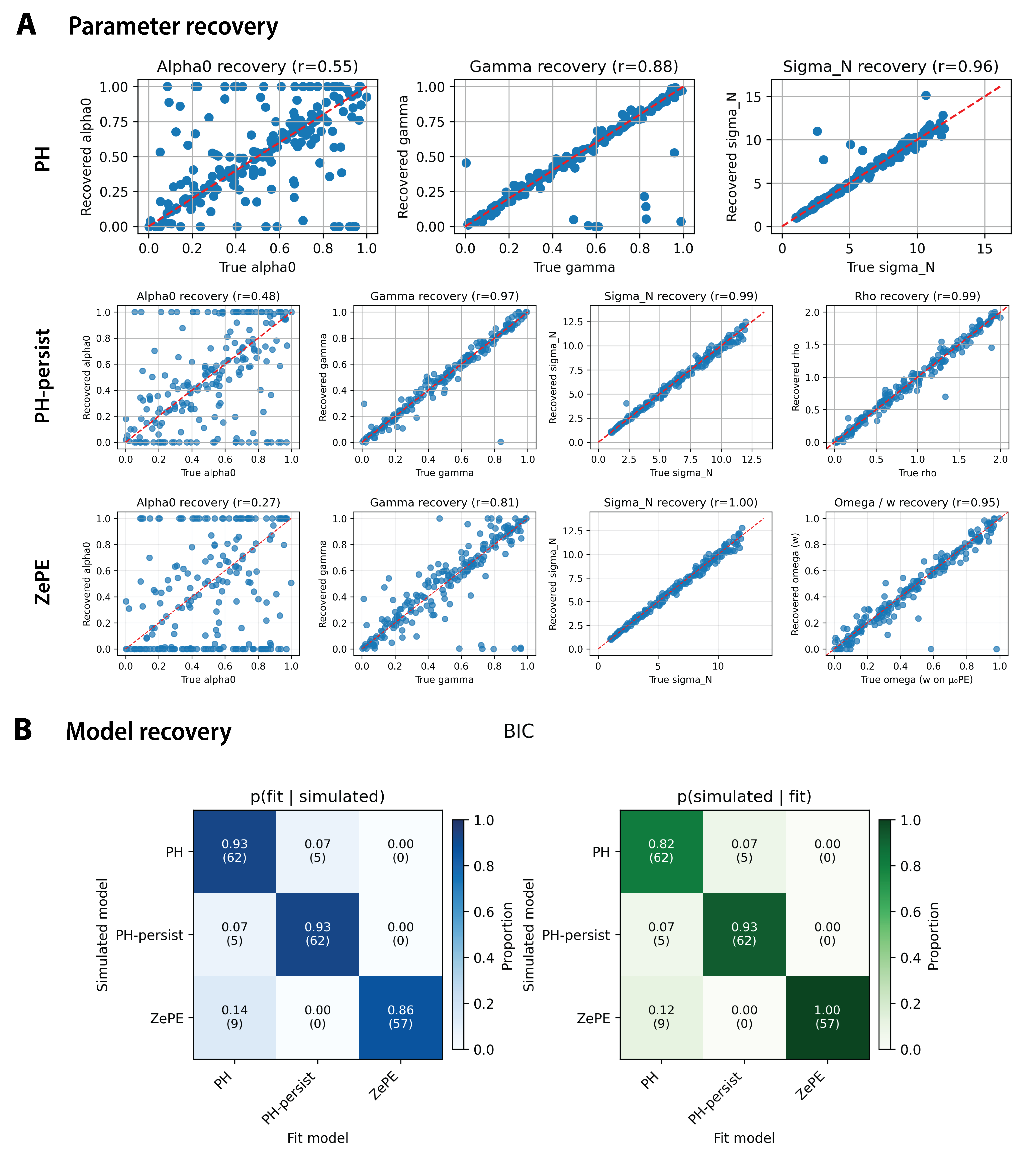

**Supplementary Figure 2.** **Parameter and model recovery.** A) Scatter plots show recovered versus true parameter values for each model (PH, PH-persist, and ZePE model). Each point represents one simulated dataset. The dashed red line indicates the identity line (perfect recovery). Pearson correlation coefficients (r) between true and recovered parameters are shown in each panel. Recovery was strongest for parameters central to each model (γ, ρ, ω, and σ), while recovery of the initial learning-rate parameter (α₀) was comparatively weaker. B) Confusion matrices showing model identifiability based on BIC. Left: probability of selecting each fitted model given the true (simulated) model. Right: probability of the true model given the selected fitted model. Values indicate proportions, with counts shown in parentheses. Diagonal elements reflect correct model identification. Overall, models were well distinguished, with high recovery accuracy and limited misclassification.

**Supplementary Table 5.**

*Task-adjusted combined results: ZePE model parameters and omega*
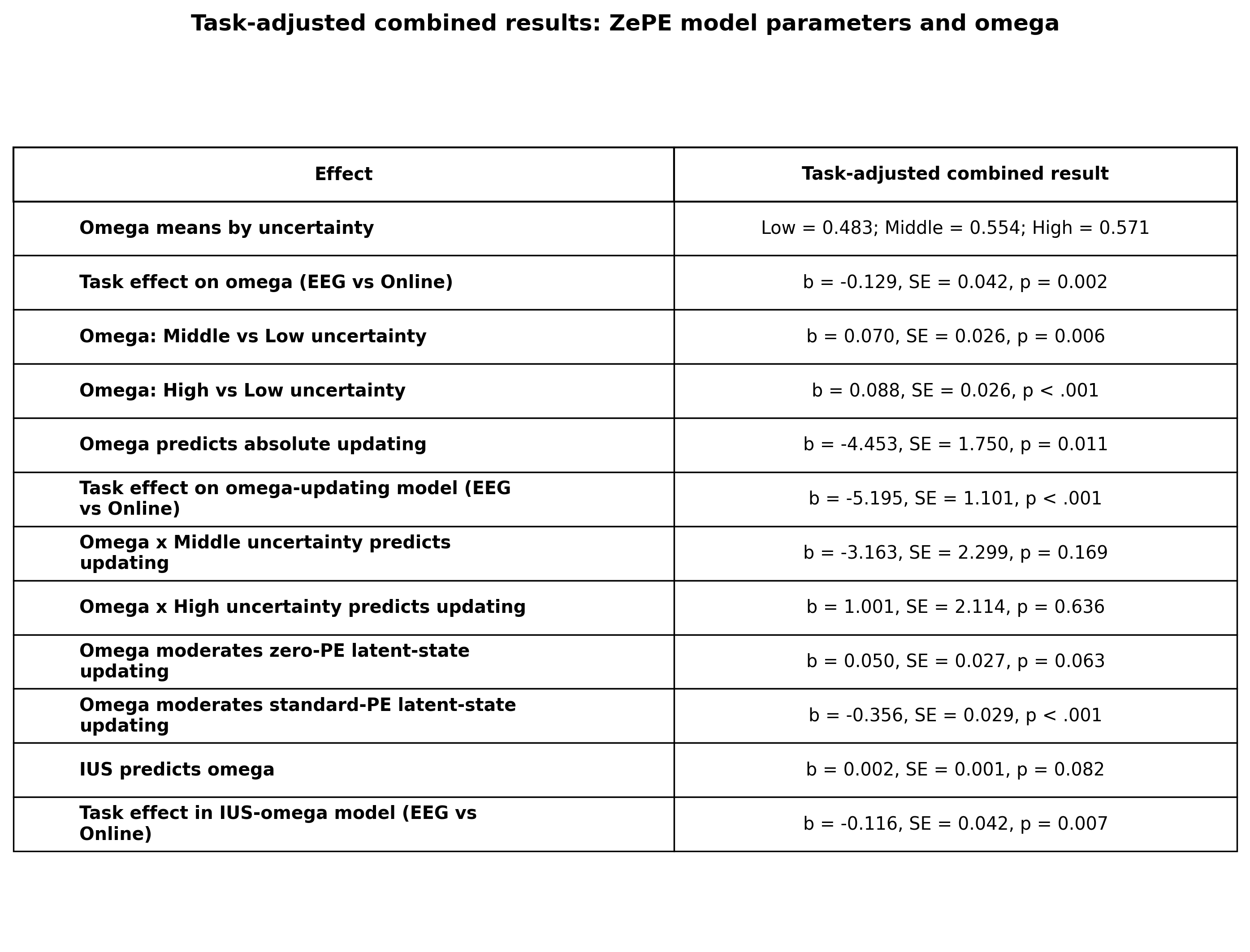

*Note*. Table shows task-adjusted combined results for ZePE model parameters and omega across the Online and EEG samples. Task was included as an additional fixed-effect regressor, with the online sample as the reference category. Omega (ω) reflects the relative weighting of the zero-prediction-error latent state. Estimates are regression coefficients with standard errors, except for omega means, which are reported by uncertainty level. Omega increased with uncertainty and was associated with subsequent updating after controlling for task, block, and trial number.

***Behavioral updating mirrors the patterns observed in simulations of ZePE***

We carried out simulations of the ZePE model in which the mixture parameter ω was systematically varied across its full range in steps of 0.33. These simulations showed that the strength of zero prediction error related updating depended on ω: larger ω values attenuated the relationship between prediction error and update magnitude, producing a shallower prediction-error–update function (Supplementary Fig. 3). Qualitatively, the behavioral data showed the same pattern: when observations were grouped into tertiles based on estimated ω values, participants with lower ω showed steeper PE-to-update relationships, whereas participants with higher ω showed shallower updating. We tested this relationship using a linear mixed-effects model with continuous ω and uncertainty level as predictors. This analysis showed that the relationship between ω and absolute prediction updates differed significantly by uncertainty level, with stronger attenuation in Middle blocks, ω × Middle: *b* = −4.31, *p* < .001, and High blocks, ω × High: *b* = −4.28, *p* < .001, relative to Low blocks. Thus, higher ω was associated with reduced behavioral updating, particularly under greater uncertainty, providing behavioral support for the interpretation of ω as capturing an attenuation of updating in the best-fitting ZePE model.

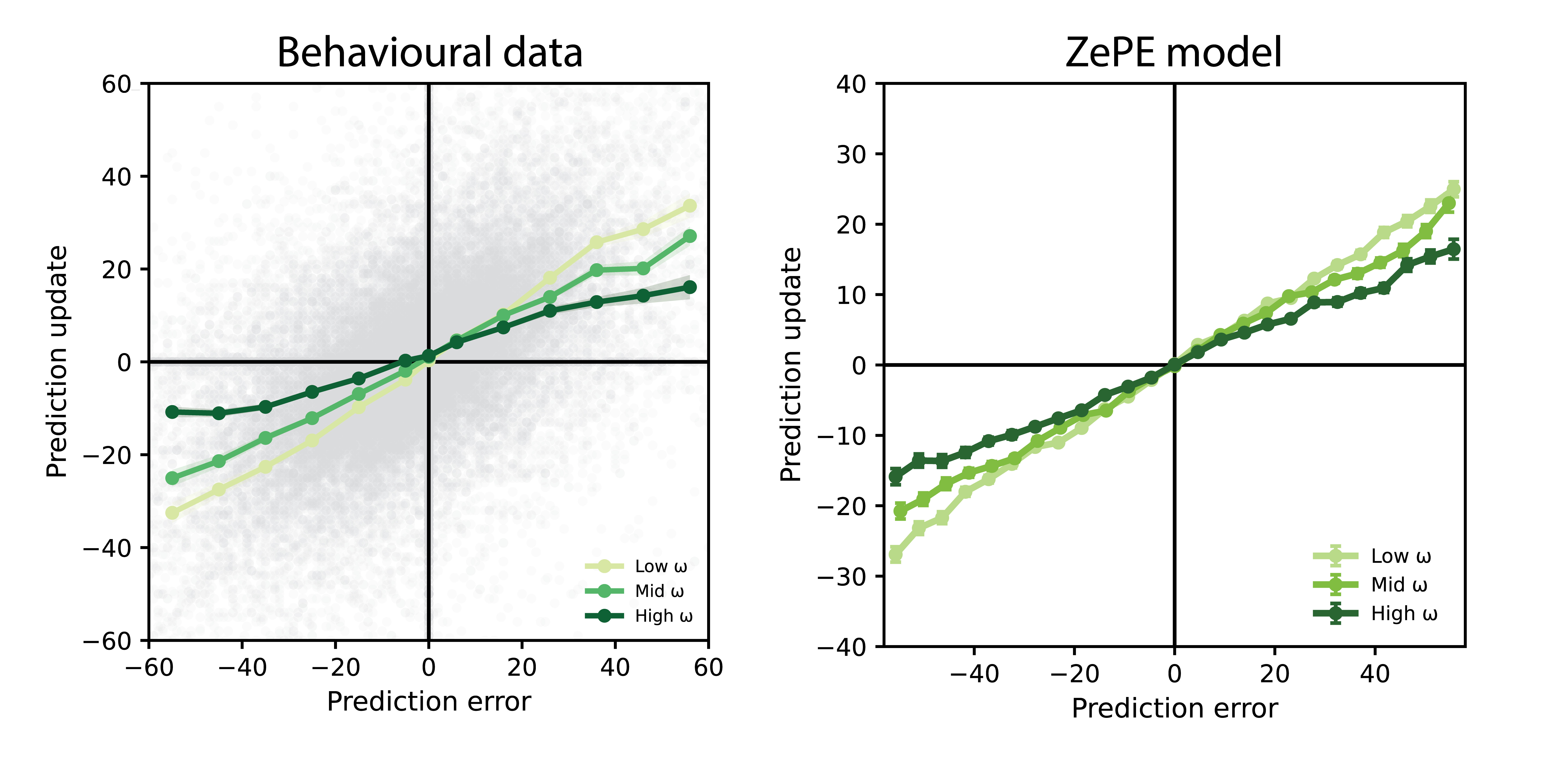

**Supplementary Figure 3.** Prediction updates from the behavioral data (left) and simulated prediction updates from the ZePE model (right), split by ω tertile. Higher ω values were associated with attenuated prediction updates. Data points depict individual participants and error bars and shaded regions denote SEM.

***EEG signals linked to (zero) prediction errors***

First, we carried out a mass-univariate regression analysis to assess EEG signals related to absolute prediction errors. To this end, we regressed trial-wise EEG activity onto absolute prediction error while controlling for uncertainty, block number, and trial number in a mass univariate model across all timepoints and electrodes. This GLM revealed two significant spatiotemporal clusters in which EEG activity scaled with absolute prediction error (cluster-based permutation test, *p* < .05; Supplementary Fig. 4A). Cluster extent is reported descriptively, as cluster-based permutation tests do not establish the precise latency or spatial location of an effect ^5^. The first cluster included time points from approximately 200–320 ms following reward onset and encompassed widespread frontocentral and parietal electrodes (*p* = .041, mean *t* = 4.01). This was followed by a later cluster including time points from approximately 420–700 ms, with a similarly broad spatial distribution (*p* = .004, mean *t* = −3.42).

To test whether the prediction error effects we found in our mass-univariate regression approach (Supplementary Fig. 4A) were specifically driven by zero prediction errors, we extended the original model to include a binary regressor coding zero-prediction-error trials (EEG ~ zero PE + |PE| + nuisance terms). This GLM revealed robust spatiotemporal clusters associated with zero prediction errors. An early positive change in voltage was observed from ~20–360 ms (*p* = .002; mean *t* = 2.9; Supplementary Fig. 4B), followed by a sustained negative effect spanning 400–1000 ms (*p* = .0005; mean *t* = −3.22), both broadly distributed across electrodes and driven by the presence of zero-prediction-error feedback. The late zero-prediction-error effect was particularly pronounced, with a sustained negative change in the amplitude peaking around ~600 ms over frontocentral electrodes (Supplementary Fig. 4B topography). This temporal profile closely overlapped with the late cluster observed in the |prediction error| model, suggesting that zero prediction errors account for the dominant late component of the reward-locked response.

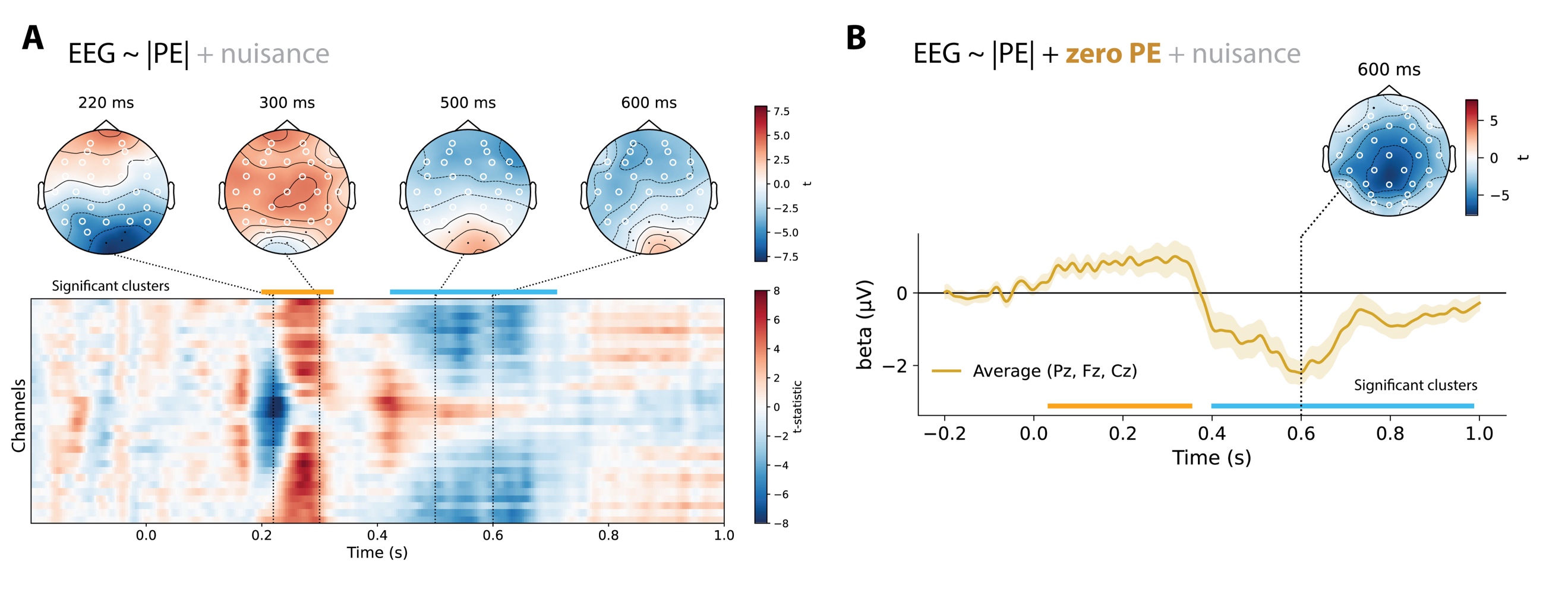

**Supplementary Figure 4. Electrophysiological signatures of (zero) prediction error.** A) Spatiotemporal regression results for absolute prediction error magnitude (|PE|). Topographies show regression coefficients at selected time points (220, 300, 500, 600 ms). The heatmap displays *t*-statistics across time (x-axis) and channels (y-axis), with significant clusters indicated above (orange: positive, blue: negative). B) Time course of regression coefficients for zero prediction errors, controlling for |PE|, averaged across frontocentral electrodes (Pz, Fz, Cz). Horizontal bars and white circles indicate spatiotemporal clusters that survived cluster-based permutation testing (*p* < .05). Error bars and shaded regions denote SEM.

**Supplementary Table 6.**

*Mixed-effects PPI model predicting trial-by-trial belief updating from residual EEG activity, latent belief-state prediction errors, and trial type.*

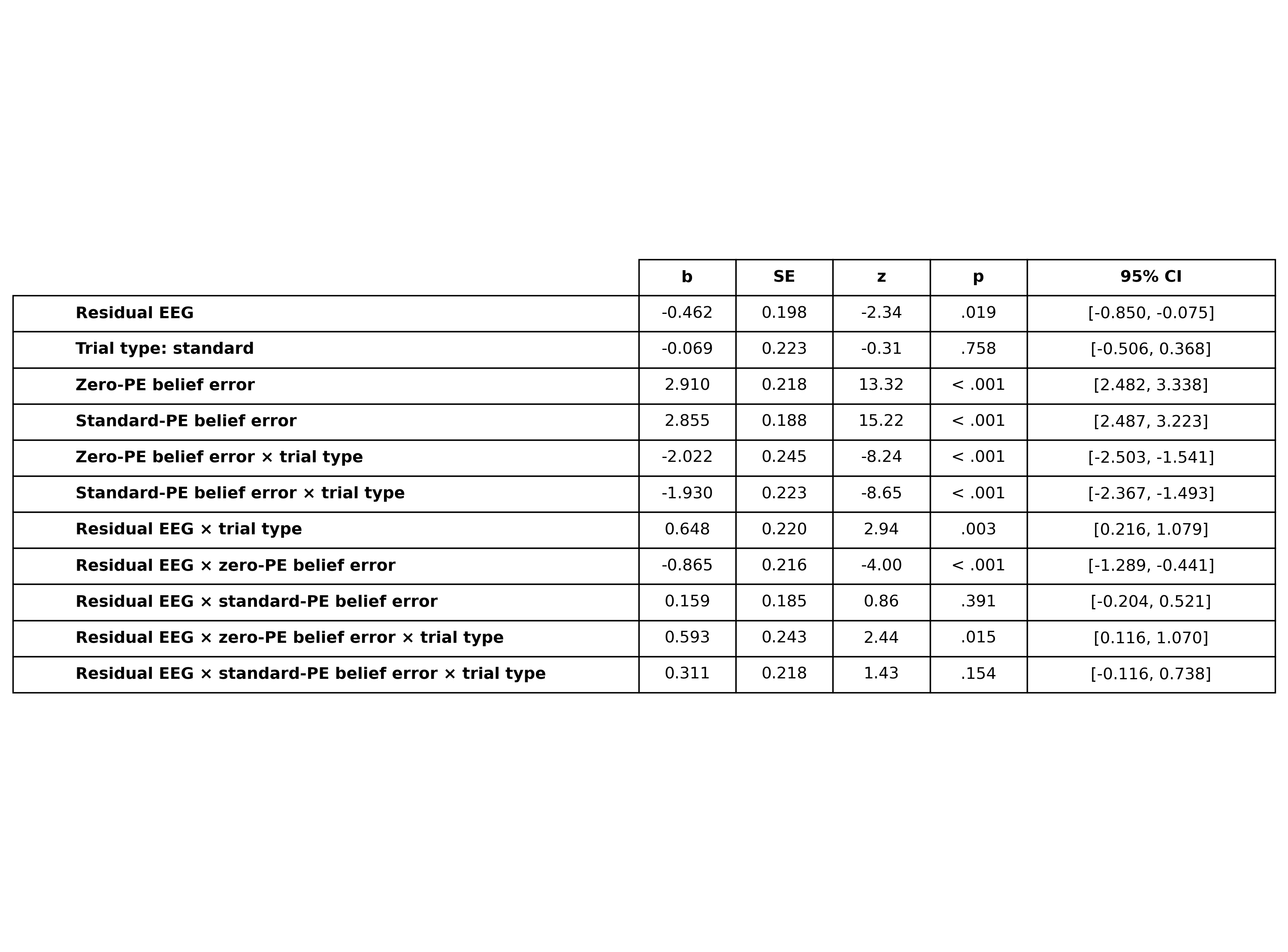

*Note.* The model tested whether residual EEG activity interacted with zero-prediction-error and standard-prediction-error belief-state signals to predict subsequent absolute prediction updating. Fixed effects shown exclude nuisance covariates (uncertainty level, block number, and trial number). Coefficients are reported with standard errors, z statistics, p values, and 95% confidence intervals.

**Supplementary Table 7.**

*Mixed-effects model predicting trial-by-trial belief updating from residual EEG activity, latent belief-state prediction errors, trial type, and ω.*

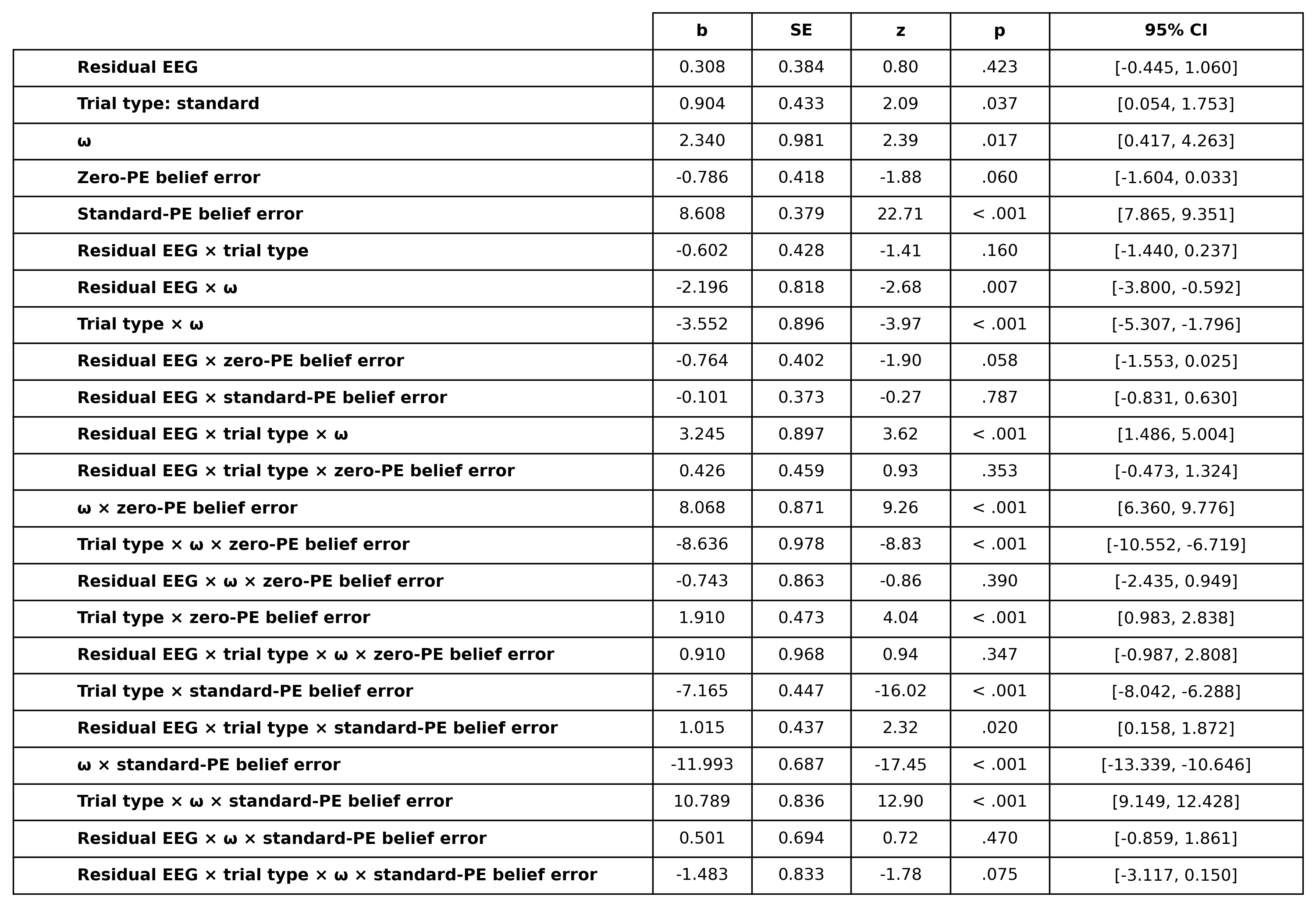

*Notes.* The model tested whether residual EEG activity predicted subsequent absolute prediction updating as a function of trial type, omega (ω), and latent belief-state prediction errors associated with zero-prediction-error and standard-prediction-error states. The model included higher-order interactions between residual EEG activity, trial type, ω, and belief-state prediction errors. Fixed effects shown exclude nuisance covariates (uncertainty level, block number, and trial number). Positive coefficients indicate larger subsequent absolute belief updates. Coefficients are reported with standard errors (SE), z statistics, p values, and 95% confidence intervals (CI). Trial type was treatment coded with zero-prediction-error trials as the reference level. ω denotes the mixture-weighting parameter from the ZePE learning model.
